# Hookworm genes encoding intestinal excreted-secreted proteins are transcriptionally upregulated in response to the host’s immune system

**DOI:** 10.1101/2025.02.01.636063

**Authors:** Erich M. Schwarz, Jason B. Noon, Jeffrey D. Chicca, Carli Garceau, Hanchen Li, Igor Antoshechkin, Vladislav Ilík, Barbora Pafčo, Amy M. Weeks, E. Jane Homan, Gary R. Ostroff, Raffi V. Aroian

**Affiliations:** Department of Molecular Biology and Genetics, Cornell University, Ithaca, NY, 14853, USA; Program in Molecular Medicine, University of Massachusetts Chan Medical School, Worcester, MA, 01605, USA; Division of Biology and Biological Engineering, California Institute of Technology, Pasadena, CA, 91125, USA; Department of Botany and Zoology, Faculty of Science, Masaryk University, Kotlářská 267/2, 611 37 Brno, Czech Republic; Institute of Vertebrate Biology, Czech Academy of Sciences, Květná 8, 603 65 Brno, Czech Republic; Department of Biochemistry, University of Wisconsin-Madison, Madison, WI, 53706, USA; ioGenetics LLC, 301 South Bedford Street, Ste.1, Madison, WI, 53703, USA

## Abstract

Hookworms are intestinal parasitic nematodes that chronically infect ∼500 million people, with reinfection common even after clearance by drugs. How infecting hookworms successfully overcome host protective mechanisms is unclear, but it may involve hookworm proteins that digest host tissues, or counteract the host’s immune system, or both. To find such proteins in the zoonotic hookworm *Ancylostoma ceylanicum*, we identified hookworm genes encoding excreted-secreted (ES) proteins, hookworm genes preferentially expressed in the hookworm intestine, and hookworm genes whose transcription is stimulated by the host immune system. We collected ES proteins from adult hookworms harvested from hamsters; mass spectrometry identified 565 *A. ceylanicum* genes encoding ES proteins. We also used RNA-seq to identify *A. ceylanicum* genes expressed both in young adults (12 days post-infection) and in intestinal and non-intestinal tissues dissected from mature adults (19 days post-infection), with hamster hosts that either had normal immune systems or were immunosuppressed by dexamethasone. In adult *A. ceylanicum*, we observed 1,670 and 1,196 genes with intestine- and non-intestine-biased expression, respectively. Comparing hookworm gene activity in normal versus immunosuppressed hosts, we observed almost no changes of gene activity in 12-day young adults or non-intestinal 19-day adult tissues. However, in intestinal 19-day adult tissues, we observed 1,951 positively immunoregulated genes (upregulated at least two-fold in normal hosts versus immunosuppressed hosts), and 137 genes that were negatively immunoregulated. Thus, immunoregulation was observed primarily in mature adult hookworm intestine directly exposed to host blood; it may include hookworm genes activated in response to the host immune system in order to neutralize the host immune system. We observed 153 ES genes showing positive immunoregulation in 19-day adult intestine; of these genes, 69 had ES gene homologs in the closely related hookworm *Ancylostoma caninum*, 24 in the human hookworm *Necator americanus*, and 24 in the more distantly related strongylid parasite *Haemonchus contortus*. Such a mixture of rapidly evolving and conserved genes could comprise virulence factors enabling infection, provide new targets for drugs or vaccines against hookworm, and aid in developing therapies for autoimmune diseases.

## Introduction

The hookworms *Necator americanus*, *Ancylostoma duodenale*, and *Ancylostoma ceylanicum* are parasitic nematodes that infect ∼500 million human beings, sickening them and lowering their economic productivity (Bethony *et al*. 2006; Traub 2013; Pullan *et al*. 2014). Hookworms are remarkably long-lived parasites: an adult hookworm can feed off its human host for up to 18 years (Beaver 1988; Gems 2000), and infection times of 1-5 years are common (Hoagland and Schad 1978). Hookworms achieve this partly by feeding on the host with enzymes optimized to digest host proteins (Ranjit *et al*. 2009), and partly by suppressing the immune systems of their hosts, which would otherwise kill or expel them quickly (Maizels *et al*. 2018). Only one drug, albendazole, is commonly used and effective enough to be useful in mass drug administration against hookworm infections (Keiser and Utzinger 2008; Loukas *et al*. 2016). However, human hookworms may be developing genetic resistance to this drug (Orr *et al*. 2019; Vlaminck *et al*. 2019; Walker *et al*. 2021), as has already happened with dog hookworms (Venkatesan *et al*. 2023). Moreover, even effective use of albendazole does not prevent endemic hookworms from reinfecting people (Jia *et al*. 2012). An anti-hookworm vaccine would be an ideal way to suppress hookworm infections (Cohen 2016), but no such vaccine exists, in part because we do not know which gene products mediating host-parasite interactions should be targeted as antigens (Zawawi and Else 2020).

It is not yet certain how hookworms (or any other parasitic nematodes) dampen the host immune system, but many possible effectors of immunomodulation exist: secreted proteases (Ranjit *et al*. 2009; Knox 2012; Pearson *et al*. 2012); secreted protease inhibitors (Manoury *et al*. 2001; Hartmann and Lucius 2003; Klotz *et al*. 2011); large gene families encoding diverse secreted proteins as antigenic decoys (Cantacessi *et al*. 2009; Tribolet *et al*. 2015); mimics of mammalian immune proteins (Loukas *et al*. 2000; Yoshida *et al*. 2012); lipid immunomodulators (Harnett *et al*. 2010; Jex *et al*. 2014; Shinoda *et al*. 2022); and secreted exosomes with anti-host miRNAs (Buck *et al*. 2014; Coakley *et al*. 2017). These mechanisms are by no means exclusive; some or all of them could be working at once. Such complexity may have evolved over 350 million years, when vertebrates first colonized the land and became vulnerable to parasitic nematodes (Anderson 1984; Clark 1994; Dsurette-Desset *et al*. 1994). Ignorance of how hookworms immunosuppress their hosts makes it challenging to devise vaccines against hookworm disease. It also hinders the possible use of hookworms as sources of new biological reagents against autoimmune diseases (Smallwood *et al*. 2021; Ryan *et al*. 2022), which appear to become more frequent with lower rates of helminth infection (Maizels 2020).

Hookworms and other parasitic helminths have long been known to excrete or secrete proteins into their hosts, and these excreted/secreted (ES) proteins have long been hypothesized, and in some cases demonstrated, to enable immunosuppression and virulence (Stirewalt 1963; Pritchard 1986; Lightowlers and Rickard 1988; Maizels *et al*. 2018; Abuzeid *et al*. 2020). ES proteins can be exported from cephalic/pharyngeal glands, intestine, or cuticle (Pritchard 1986; Zhan *et al*. 2003; Huang *et al*. 2020). Genes encoding ES proteins have been identified in the major human hookworm *N. americanus* (Logan *et al*. 2020), the dog hookworm *Ancylostoma caninum* (Morante *et al*. 2017), and five strongylid parasitic nematodes closely related to hookworm: *Angiostrongylus vasorum* (Gillis-Germitsch *et al*. 2021), *Haemonchus contortus* (Wang *et al*. 2019), *Heligmosomoides bakeri* (Hewitson *et al*. 2011; Moreno *et al*. 2011; Hewitson *et al*. 2013), *Nippostrongylus brasiliensis* (Sotillo *et al*. 2014), and *Ostertagia ostertagi* (Price *et al*. 2024) (Figure 1). Until recently (Uzoechi *et al*. 2023), ES proteins have not been identified for the zoonotic human hookworm *A. ceylanicum*, which is an important zoonotic pathogen of humans and companion animals and which can complete its lifecycle in hamsters, making it a key hookworm species for laboratory studies (Garside and Behnke 1989; Traub 2013; Stracke *et al*. 2020; Colella *et al*. 2021).

**Figure 1.**
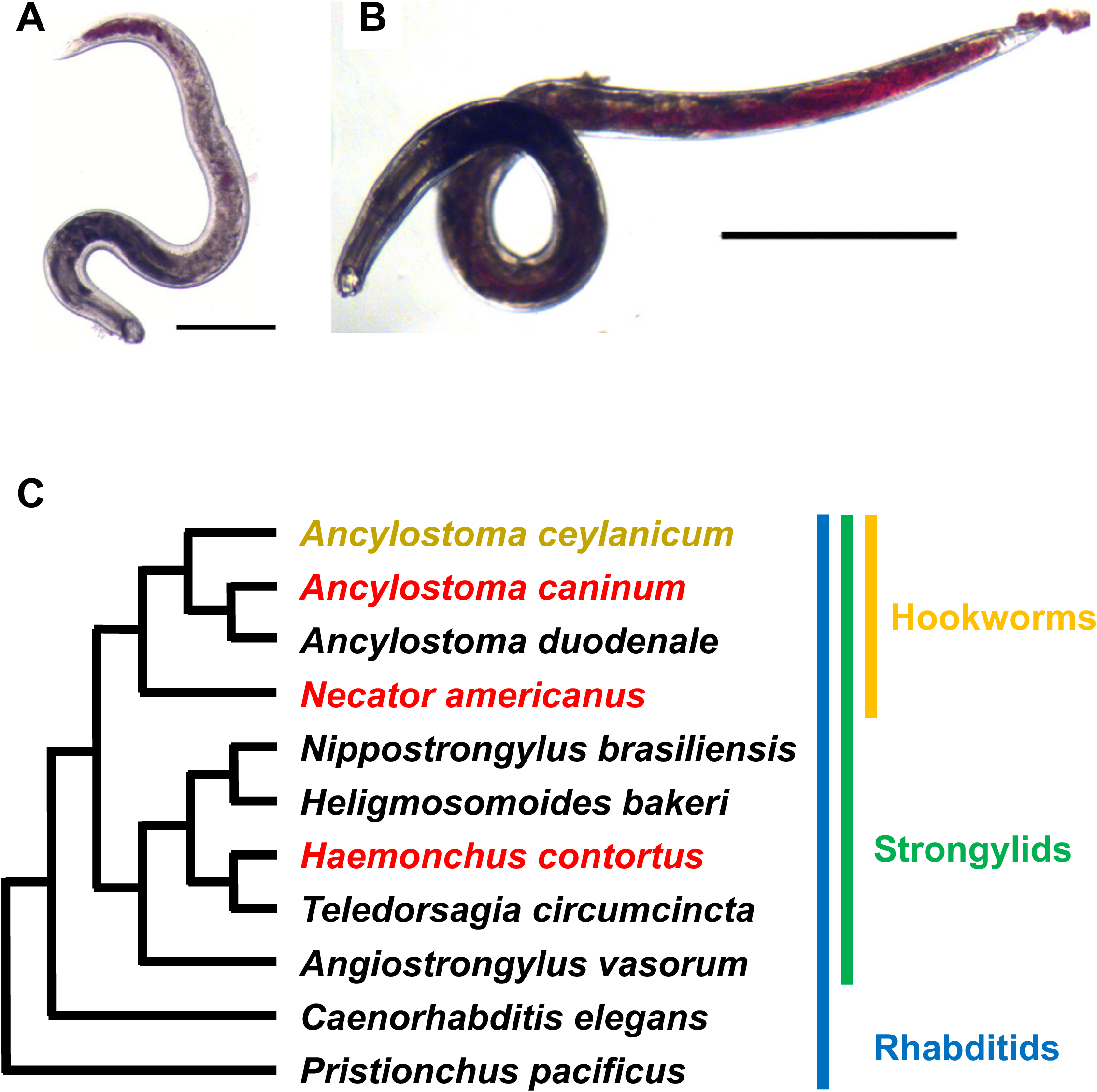
Morphology and evolution of the hookworm *Ancylostoma ceylanicum*. (A) Female of *A. ceylanicum* by 12 days after infection of a hamster host; scale bar, 500 μm. At this stage of development, the hookworms are young adults that have only just begun blood feeding, with mature males and a few gravid females, and with little or no egg laying. (B) Male of *A. ceylanicum* by 19 days after infection of a hamster host; scale bar, 1 mm. At this stage of development, the hookworms are fully mature adults that have been blood-feeding for at least a week, have mated, and have begun extensive egg laying that can last for weeks in a hamster host. Both photographs are reproduced from Schwarz *et al*. (Schwarz *et al*. 2015). (C) Evolutionary relationships of the hookworm *A. ceylanicum* to some other nematodes discussed in this paper. This phylogeny links *A. ceylanicum* to other *Ancylostoma* and *Necator* hookworms, to more distantly related strongylid parasitic nematodes, and to two well-studied free-living rhabditid nematodes (*C. elegans* and *Pristionchus pacificus*). Phylogenetic data are taken from Coghlan *et al*., van Megen *et al*., and Xie *et al*. (van Megen *et al*. 2009; Xie *et al*. 2017; Coghlan *et al*. 2019). *A. ceylanicum* is highlighted in gold; three parasitic species (*A. caninum*, *N. americanus*, and *H. contortus*) whose ES genes were compared to those of *A. ceylanicum* in this paper are highlighted in red. Hookworms form a single clade (marked with a yellow bar); they are a subset of a larger clade of strongylid nematodes (marked with a green bar) which, like hookworms, are parasitic. Strongylids are, in turn, a subset of rhabditid nematodes (marked with a blue bar); this clade encompasses both parasitic nematodes and the free-living nematodes *C. elegans* and *P. pacificus*. Notably, *A. ceylanicum* and other strongylid parasites are more closely related to *C. elegans* than to *P. pacificus* despite having highly divergent parasitic life cycles.

To find *A. ceylanicum* genes whose products may enable parasitism, we have identified *A. ceylanicum* ES proteins, while also using RNA-seq to identify *A. ceylanicum* genes whose products interact with the host either through intestinally-biased expression or through upregulation in response to a functioning host immune system. For the latter set of genes, we hypothesized that the parasite transcriptionally activates genes in response to the host immune system in order to neutralize the host immune response. Previously, we found that changes of *A. ceylanicum* gene activity during infection *in vivo* are much more extensive than changes seen during simulated infection *in vitro* (Schwarz *et al*. 2015). Here we build on that observation by correlating ES genes with intestinal and immunoregulated genes. We identify genes encoding immunoregulated intestinal ES proteins that may be important for virulence or immunosuppression, that are new targets for drugs or vaccines against hookworm, and that may aid in the development of therapies against autoimmune diseases.

## Methods

### Sample procurement, preparation and storage

Cultures, infections, and collections of *A. ceylanicum* followed published methods (Hu *et al*. 2012; Schwarz *et al*. 2015). For each *A. ceylanicum* infection in support of RNA-seq, we purchased Syrian golden hamsters (*Mesocricetus auratus*) of the HsdHan:AURA strain at 3-4 weeks of age. Hamsters were provided with food and water *ad libitum*. For immunosuppression experiments, we immunosuppressed half of each set of hamsters by injecting them with dexamethasone (3 mg/kg) twice per week throughout the duration of the experiment (Tritten *et al*. 2012); the other half were given mock injections. After the first two injections (one week), both immunocompetent and immunocompromised hamsters were infected at approximately four to five weeks of age with *A. ceylanicum*. Infections were allowed to progress for 12 or 19 days, after which we euthanized the hamsters and collected hookworms from small intestines dissected from the hamsters. Dissected hamster intestines were put into Hank’s Balanced Salt Solution (pre-warmed to 37°C); worms were picked quickly from the dissected tissues by hand. Collected worms were snap-frozen in liquid nitrogen and stored at −80°C until use. For *A. ceylanicum* infections in support of ES protein mass spectrometry, we followed a similar protocol but without injections, with two collections of *A. ceylanicum* at 20 days after infection, and without snap-freezing of dissected hookworms. Stages of *A. ceylanicum* selected for RNA-seq or proteomics are based on previously described stages of growth in golden hamsters (Ray *et al*. 1972). For hookworm intestinal-specific studies, from each triplicate set of 19-day post-infection hookworms isolated from hamsters as above, we dissected both hookworm intestinal and hookworm non-intestinal tissue, and extracted RNA from the dissected tissues; for the triplicates of smaller 12-day post-infection young adult worms, such dissection was not possible, so we extracted RNA from whole 12-day young adult worms.

All animal experiments were carried out under protocols approved by the University of Massachusetts Chan Medical School. All housing and care of laboratory animals used in this study conformed to the NIH Guide for the Care and Use of Laboratory Animals in Research (18-F22) and all requirements and all regulations issued by the USDA, including regulations implementing the Animal Welfare Act (P.L. 89-544) as amended (18-F23).

### RNA harvesting and sequencing

Total *A. ceylanicum* RNA was extracted from either whole worms or from dissected tissues as in Romeo and Lemaitre (Romeo and Lemaitre 2008; Schwarz *et al*. 2015). RNA-seq was done largely as in Srinavasan *et al*. (Srinivasan *et al*. 2013). RNA-seq libraries were built with Illumina’s TruSeq RNA Sample Prep Kit v2 executed according to manufacturer’s instructions, using 1 µg of total RNA for each sample. RNA-seq libraries were sequenced in single-end mode with read lengths of 50 nt (for all 19-day data) or 100 nt (for all 12-day data). Newly generated *A. ceylanicum* RNA-seq libraries are listed in Supplementary Table S1.

### Protein harvesting

After 20 days of infection, *A. ceylanicum* hookworms were isolated from their hamster hosts and cultured in liquid culture medium for up to 3 days at 37°C in 5% CO_2_. Liquid culture medium consisted of RPMI Medium 1640 (Gibco, Cat#11835-030) supplemented with 25 mM pH 7.2 HEPES buffer, 10 µg/ml amphotericin B (Gibco, Cat# 15290-026), and 100 U/ml penicillin/streptomycin (Gibco, Cat# 1570-063), with medium sterilized with a 0.22 µm filter. Fetal bovine serum was omitted from the medium to eliminate any added protein when quantifying ES proteins by BCA assay and to eliminate contamination for proteomics. After the first 24 hours, proteins excreted or secreted from the hookworms were collected by aspirating media only. The aspirated media were centrifuged to remove debris (1500 gs for 20 minutes, at 4°C); the resulting supernatant was concentrated 10-fold using a 3kD Amicon ultra-centrifugal filter (4000 gs for 45 minutes, at 4°C). After this first 10-fold concentration, PBS was added to match the initial volume, after which the media were reconcentrated by again being centrifuged (4000 gs for 45 minutes, at 4°C); we repeated this twice, for a total of three PBS rinses and reconcentrations. After the third PBS rinse/reconcentration, total protein was quantified using a Pierce BCA Protein Assay kit. Purified ES proteins were then frozen and stored at −80°C.

### General computation

Where possible, we used mamba to install and run version-controlled software environments from bioconda (Gruning *et al*. 2018). For reformatting or parsing of computational results, we used Perl scripts either developed for general use or custom-coded for a given analysis. All such Perl scripts (named below with italics and the suffix “*.pl*”) were archived on GitHub (https://github.com/schwarzem/ems_perl). Internet sources (URLs) for other software are listed in Supplementary Table S2.

### Nematode genomes, transcriptomes, coding sequences, and proteomes

For transcriptomic or proteomic analyses, published genome sequences, coding sequences, proteomes, and gene annotations of relevant nematodes were downloaded from WormBase (Davis *et al*. 2022) or ParaSite (Howe *et al*. 2017; Lee *et al*. 2017) (Supplementary Table S3). Published RNA-seq data of *A. ceylanicum* (Schwarz *et al*. 2015; Wei *et al*. 2016; Bernot *et al*. 2020) and *H. contortus* (Laing *et al*. 2013) were downloaded from the European Nucleotide Archive (Supplementary Table S4). Alternative gene predictions for *A. ceylanicum* recently published by Uzoechi *et al*. (Uzoechi *et al*. 2023) were obtained as a Generic Feature Format Version 3 (GFF3) annotation file (https://github.com/The-Sequence-Ontology/Specifications/blob/master/gff3.md) from Young-Jun Choi that we have archived at https://osf.io/dxfsb. We extracted protein sequences from this GFF3 via gffread 0.12.7 (Pertea and Pertea 2020) with the arguments ‘*-g [input genome sequence FASTA file] -o /dev/null -C --sort-alpha -- keep-genes -P -V -H -l 93 -y [output protein FASTA file] [input gene prediction GFF3 file]*’.

Heligmosomoides *species nomenclature*. Recent genomic analysis has shown that the gastrointestinal parasitic nematode *Heligmosomoides* has two distinct species: *H. bakeri* and *H. polygyrus* (Stevens *et al*. 2023). Although laboratory strains of *Heligmosomoides* have often been described as *H. polygyrus* (Maizels *et al*. 2011), it now appears likely that many or all of these strains have actually been *H. bakeri*. Thus, following previous suggestions for revised nomenclature (Behnke and Harris 2010), we refer exclusively to *H. bakeri* even when citing published work that was nominally done with *H. polygyrus*.

### RNA-seq subsampling

We found that published *A. ceylanicum* RNA-seq data (Supplementary Table S4) were too extensive for gene predictions by BRAKER2 because of memory limitations. To make these data usable by BRAKER2, we selected a representative subset of them with khmer (Zhang *et al*. 2014; Crusoe *et al*. 2015). Because khmer requires “#/1” and “#/2” suffixes for paired-end reads, we retrofitted paired-end RNA-seq reads lacking such suffixes with *retroname_fastq_reads.pl*. We ran *normalize-by-median.py* from khmer on all data (paired and unpaired) twice, with the arguments *’-k 31 -C 100 -M 100G*’, and *’-k 31 -C 30 -M 100G*’; we then ran *filter-abund.py* from khmer on paired-end data with the arguments ‘*--variable-coverage -C 2 [k-mer hash] [paired-end reads]*’. We sorted khmer-filtered data into interleaved paired- and unpaired-end read files with *paired_vs_unp_fastq.or.a.pl* with the arguments ‘*--r1 “#0\/1” --r2 “#0\/2“*’.

### Reprediction of protein-coding genes

To repredict protein-coding genes in *A. ceylanicum*, we ran *braker.pl* in BRAKER2 2.1.6 (BrŮna *et al*. 2021) on our published repeat-softmasked *A. ceylanicum* genome assembly (Schwarz *et al*. 2015), with the arguments ‘*--genome [genome assembly FASTA] -- prot_seq [collected proteomes FASTA] --bam [sorted mapped khmer-filtered paired-end read BAM alignment] --etpmode --softmasking --cores 48 --gff3*’. BRAKER2 requires an input genome sequence to have its FASTA header lines previously stripped of comments; we did this with *uncomment_FASTA_headers.pl*. Running BRAKER2 also required us to install: AUGUSTUS 3.4.0 (Stanke *et al*. 2008); BamTools 2.5.2 (Barnett *et al*. 2011); cdbfasta 0.99 (Pertea *et al*. 2003); DIAMOND 2.0.9 (Buchfink *et al*. 2021); GeneMark-ES/ET 4.68 (Lomsadze *et al*. 2014); and SAMtools 1.12 (Danecek *et al*. 2021). To guide BRAKER2 gene predictions, we used our khmer-subsampled paired-end subset of published *A. ceylanicum* RNA-seq data, along with predicted proteomes from the related nematodes *Caenorhabditis elegans* (Davis *et al*. 2022), *H. contortus* (Doyle *et al*. 2020), and *N. americanus* (Logan *et al*. 2020). After running BRAKER2, we renamed gene, transcript, and exon names in the GFF3 prediction file with a modified version of *updateBRAKERGff.py* (https://github.com/Gaius-Augustus/BRAKER/issues/416) and Perl one-line commands. To extract coding DNAs (CDS DNAs) and protein sequences from the renamed GFF3, we used gffread 0.12.7 (Pertea and Pertea 2020) as above, with the additional argument ‘*-x [output CDS DNA file]*’.

This gave us a new gene set (“v2.0”) that was generally superior (in completeness as assayed by BUSCO, and nonfragmentation as assayed by *count_fasta_residues.pl*) to our original 2015 gene set (“v1.0”). However, we observed that some v1.0 genes encoded ES proteins (as determined by mass spectrophotometric mapping) but had no genomic overlap with v2.0 genes. To ensure no valid ES genes would be overlooked, we used BEDtools 2.30.0 (Quinlan and Hall 2010) to identify v1.0 genes whose protein-coding exon sequences (CDSes) had no genomic overlap with v2.0 CDSes, as follows. We used *get_gff3_gene_subset.pl* to remove GFF3 v2.0 annotations that could not be translated by gffread, and then used Perl to extract CDS annotation lines from the GFF3 files of published v1.0 and gffread-translatable v2.0 gene predictions. We reformatted the CDS-subset files from GFF3 to BED with *gff2bed* in BEDOPS 2.4.41 (Neph *et al*. 2012). We identified overlapping coordinates of the CDS-only BED annotation files for v1.0 and v2.0 genes with *intersect* from BEDtools 2.30.0 (Quinlan and Hall 2010), using the argument ‘*-loj*’, and used Perl to extract a list of v1.0 genes having no CDS overlaps with v2.0 genes. Given this list of non-overlapping v1.0 genes, we used *extract_fasta_subset.pl* to extract their encoded CDS DNAs and proteins as subsets from the full published v1.0 CDS DNA and protein sets, and added these CDS DNA and protein subsets to our previous generated v2.0 CDS DNA and protein sequences. This gave us our final hybrid prediction set (“v2.1”) of CDS DNAs and proteins on which all further analyses in this paper were performed. The v2.1 CDS DNAs and proteome have been archived at https://osf.io/dxfsb.

We generated a v2.1 GFF3 as follows. From the published v1.0 GFF3, we extracted GFF3 annotations for these non-overlapping v1.0 genes first by running *extract_parasite_GFF3_subset.pl* and then by selecting ‘*WormBase_imported*’ annotation lines with Perl. We reformatted the resulting v1.0 non-overlapping GFF3 with AGAT 1.1.0 (Dainat 2023). Likewise, we reformatted the gffread-translatable v2.0 GFF3 with AGAT. We then merged the two AGAT-reformatted GFF3s to produce a single v2.1 GFF3 with uniform formatting, which we have archived at https://osf.io/dxfsb.

### Assessing quality of protein-coding genes

We determined general statistics of protein-coding gene products with *count_fasta_residues.pl* using the arguments ‘*-e -t prot*’; we obtained gene counts from maximum-isoform proteome subsets generated with *get_largest_isoforms.pl* using the arguments ‘*-t parasite*’ or *’-t maker*’. We determined and compared the completeness of predicted *A. ceylanicum* protein-coding gene sets with BUSCO 5.2.2 using the arguments ‘*--lineage_dataset nematoda_odb10 --mode proteins*’ (Waterhouse *et al*. 2018), which tested a given proteome against 3,131 highly conserved single-gene orthologs in nematodes.

### Gene reannotation

For the protein products of our v2.1 *A. ceylanicum* gene set, we predicted both N-terminal signal sequences and transmembrane alpha-helical anchors with Phobius 1.01 (KÄll *et al*. 2004), reformatting results with *tabulate_phobius_hits.pl*. We predicted coiled-coil domains with Ncoils 2002.08.22 (Lupas 1996), reformatting results with *tabulate_ncoils_x.fa.pl*. We predicted low-complexity regions with PSEG 1999.06.10 (Wootton 1994) using the argument ‘*-l*’ and reformatting results with *summarize_psegs.pl*. We identified protein domains with Pfam 35.0 (Mistry *et al*. 2021) database with *hmmscan* in HMMER 3.3.2 (Eddy 2009; Finn *et al*. 2016), using the arguments ‘*--cut_ga*’ to impose family-specific significance thresholds, ‘*-o /dev/null*’ to discard text outputs, and ‘*--tblout*’ to export tabular outputs; Pfam results were reformatted with *pfam_hmmscan2annot.pl*. We also identified protein domains with *interproscan.sh* in InterProScan 5.57-90.0 (Paysan-Lafosse *et al*. 2023) using the arguments ‘*-dp -iprlookup -goterms*’, and reformatting results with *tabulate_iprscan_tsv.pl*. We generated Gene Ontology (GO) terms (Ashburner *et al*. 2000; Carbon *et al*. 2021), EggNOG descriptions (Hernandez-Plaza *et al*. 2023), and KEGG codes (Kanehisa *et al*. 2023) with EnTAP 0.10.7-beta (Hart *et al*. 2020) using the argument ‘*--runP*’, and using selected UniProt (UniProt 2023) and RefSeq (O’Leary *et al*. 2016) proteome databases from highly GO-annotated model organisms (Supplementary Table S3); proteome databases were generated with *makedb* from DIAMOND 0.9.9 (Buchfink *et al*. 2021), itself bundled with EnTAP; annotations from EnTAP were reformatted from protein to gene annotation tables with *cds2gene_EnTAP_annot.pl* and *cds2gene_annot.pl*. We identified genes encoding possible antimicrobial peptides (AMP) by mapping predictions by Irvine *et al*. (Irvine *et al*. 2023) from v1.0 to v2.1 of our gene predictions. We identified orthologies between *A. ceylanicum* and other nematodes with OrthoFinder 2.5.4 (Emms and Kelly 2019) using the arguments ‘*-S diamond_ultra_sens -og*’; results were reformatted with *prot2gene_ofind.pl* and *genes2omcls.pl*. For all analyses except OrthoFinder, full proteomes were used; for OrthoFinder, we used maximum-isoform proteome subsets generated with *get_largest_isoforms.pl*. An overall annotation table was constructed from both these and RNA-seq annotations (below) with *add_tab_annots.pl*.

### Gene annotation for related parasitic nematodes

To make consistent comparisons of ES genes (or other categories of genes) between *A. ceylanicum* and other species, we annotated the protein-coding genes for three related parasites (*A. caninum*, *H. contortus*, and *N. americanus*) using the same methods as for *A. ceylanicum* above; the analyzed proteomes are listed in Supplementary Table S3.

### Selection of ES gene sets from related parasitic nematodes for comparative analysis

We selected three published ES gene sets from *A. caninum*, *H. contortus*, and *N. americanus* for comparative analysis by the following criteria. They had to have either gene identification numbers (IDs) from public genome assemblies, or, if they did not have gene IDs from valid genomes, they needed to at least have expressed sequence tag (EST) IDs for which the original sequence data were publicly available so that they could be mapped onto genomes (and thus to modern genes with proper IDs). This requirement disqualified previously published ES gene sets for *H. bakeri* and *N. brasiliensis*. Second, the gene IDs needed to be correctly mapped from protein mass spectrometric data of one species onto genes of the same species. Remarkably, this requirement disqualified the ES genes from *Angiostrongylus*, which were generated from *A. vasorum* protein mass spectrometry data but were mapped onto genes of *A. cantonensis* and *A. costaricensis*. Third, the ES genes needed to be from parasitic nematodes that had evolved parasitism in common with *A. ceylanicum* hookworms. This criterion disqualified ES genes from the giant roundworm *Ascaris suum* or the whipworm *Trichuris muris*, which evolved parasitism independently of hookworms and other strongylids.

### RNA-seq data

For *A. ceylanicum*, we generated biologically triplicated RNA-seq data for each biological condition. We also analyzed published RNA-seq data for *A. ceylanicum* (Schwarz *et al*. 2015; Wei *et al*. 2016; Bernot *et al*. 2020) and *H. contortus* (Laing *et al*. 2013), listed in Supplementary Table S4. Before analysis, we Chastity-filtered new *A. ceylanicum* RNA-seq reads with *quality_trim_fastq.pl* using the arguments *’-q 33 -m 50’*. We then quality-filtered and adaptor-trimmed new *A. ceylanicum* RNA-seq reads with fastp 0.20.0 (Chen *et al*. 2018) using the arguments ‘--*dont_overwrite --detect_adapter_for_pe --n_base_limit 0 --length_required X*’, with X = 50 for new *A. ceylanicum* data. We quality-filtered and adaptor-trimmed previously published *A. ceylanicum* and *H. contortus* RNA-seq reads with fastp 0.20.0 using the arguments ‘*--dont_overwrite --n_base_limit 0 --length_required X*’ with X = 50 for *A. ceylanicum*.

### RNA-seq expression values and significances

We quantified gene expression from RNA-seq data sets with Salmon 1.9.0, generating expression values in transcripts per million (TPM) and estimating mapped read counts per gene (Patro *et al*. 2017). To prevent spurious mappings of RNA-seq reads, we used full selective alignment to a “gentrome” (a CDS DNA set, treated as a target for real mappings, combined with its genome, treated as a decoy for spurious mappings), followed by quantification using Salmon in non-alignment mode (Srivastava *et al*. 2020; Srivastava *et al*. 2021). For Salmon’s *index* program, we used the arguments ‘*--no-version-check --keepDuplicates -t* [gentrome sequence] *-d* [decoy list]’; for Salmon’s *quant* program, we used the arguments ‘*--no-version-check --seqBias --gcBias -- posBias --libType A --geneMap* [transcript-to-gene table]’, with ‘*--unmatedReads*’ used for single-end data. Results from *quant.genes.sf* output files were reformatted with *make_salmon_TPM_slices.pl*.

### Heatmapping of RNA-seq data

To visualize and cluster gene expression values for RNA-seq replicates, we converted expression values of 0 TPM to empirical pseudozeros with *assort_tpms.pl*, with each replicate’s empirical pseudozero being defined as the lowest non-zero TPM expression value observed for all genes within that replicate. This allowed logarithmic transformation of zero expression values while defining these values in a replicate-specific way. We then converted the expression values of replicates to log_10_(TPM) scores with *log10_tsv_numbers.pl*. Finally, we heatmapped and dendrogram-clustered all replicates with ComplexHeatmap 2.18.0 (Gu 2022) using default parameters. Heatmapping showed individual non-dexamethasone (nonDEX) and dexamethasone (DEX) intestinal RNA-seq replicates that anomalously clustered with two DEX and nonDEX intestinal RNA-seq replicates, respectively (Supplementary Figure 1); these anomalous replicates were thus not used for differential gene expression analysis.

### Differential gene expression analysis

Having generated RNA-seq readcounts, we used the exactTest function of edgeR 3.36.0 (Robinson *et al*. 2010) to compute log_2_ fold-changes (log2FC) and false discovery rate (FDR) significance values (Noble 2009), to identify statistically significant changes of gene expression between pairs of biological conditions in our RNA-seq data. Dispersions were computed with edgeR’s estimateDisp function using the argument ‘*robust=TRUE*’. For this analysis, we used all non-anomalous replicates of our *A. ceylanicum* RNA-seq data (Supplementary Table S1) along with published *A. ceylanicum* RNA-seq data from whole fourth-stage larval (L4) males, whole adult males, whole adult females, and adult male intestine (Supplementary Table S4). To increase the statistical signal for differentially expressed genes, we used *filter_minimum_readcounts.pl* with the argument ‘*10*’ to remove genes which failed to achieve 10 mapped reads in any biological replicate before submitting readcounts to edgeR. An edgeR R script (Supplementary File S1) was generated with *make_edgeR_classic_scripts.pl*, and run in batch mode with R 4.1.3. In summarizing edgeR results, we defined a gene as being differentially expressed (e.g., with tissue-, sex-, or condition-biased expression) if edgeR scored it as being ≥2-fold more strongly or weakly expressed in one condition than another (either log2FC ≥ 1, or log2FC ≤ −1) with an FDR of ≤ 0.01. All other genes with intermediate expression ratios between two conditions (−1 < log2FC < 1), or with expression ratios that were determined with lower significance (FDR > 0.01) were not classified as having differential gene expression. We used edgeR’s exactTest with these criteria because it is well-adapted for analyzing small numbers of biological replicates (Schurch *et al*. 2016).

### RNA-seq and differential gene expression analysis for H. contortus

These were performed with published RNA-seq data for dissected adult female intestine and whole adult female bodies (Supplementary Table S4) (Laing *et al*. 2013) and with predicted genes from the published chromosomal reference genome (Supplementary Table S3) (Doyle *et al*. 2020), by the same methods as used for *A. ceylanicum*.

### Constructing a protein set for mass spectrophotometric analysis

To combine our newly predicted *A. ceylanicum* proteins with alternative predictions by Coghlan *et al*. (Coghlan *et al*. 2019), we ran cd-hit-2d from CD-HIT 4.8.1 (Li and Godzik 2006) with the arguments ‘*-i [v2.1 predicted proteome] -i2 [alternative published proteome by Coghlan et al.] -o [distinct Coghlan protein sequences] -d 100 -c 1.0 -M 0 -T 1 -l 5 -s 0.0 -aL 0.0 -aS 1.0*’. We then joined the v2.1 proteome with distinct Coghlan sequences to yield a nonredundant *A. ceylanicum* protein set for LC-MS/MS analysis.

### Liquid chromatography-tandem mass spectrometry (LC-MS/MS) analysis

Peptides were analyzed with an Orbitrap Exploris 480 hybrid quadrupole-Orbitrap mass spectrometer coupled to an UltiMate 3000 RSLCnano liquid chromatography system (ThermoFisher Scientific). Peptides were loaded onto an Acclaim PepMap RSLC column (75 µm × 15 cm, 2 µm particle size, 100 Å pore size, ThermoFisher Scientific) over 15 min in 97% mobile phase A (0.1% formic acid) and 3% mobile phase B (0.1% formic acid, 80% acetonitrile) at 0.5 µL/min. Peptides were eluted at 0.3 µL/min using a linear gradient from 3% mobile phase B to 50% mobile phase B over 120 min. Peptides were electrosprayed through a nanospray emitter tip connected to the column by applying 2000 V through the ion source’s DirectJunction adapter. Full MS scans were performed at a resolution of 60,000 at 200 m/z over a range of 300-1,200 m/z with an AGC target of 300% and the maximum injection time set to ‘auto’. The top 20 most abundant precursors with a charge state of 2-6 were selected for MS/MS analysis with an isolation window of 1.4 m/z and a precursor intensity threshold of 5 × 103. A dynamic exclusion window of 20 s with a precursor mass tolerance of ± 10 ppm was used. MS/MS scans were performed using HCD fragmentation using a normalized collision energy of 30% and a resolution of 15,000 with a fixed first mass of 110 m/z. The AGC target was set to ‘standard’ with a maximum injection time of 22 ms.

### Mass spectrometry data analysis

Thermo RAW files were searched against our nonredundant *A. ceylanicum* protein set using the SEQUEST algorithm in Proteome Discoverer 2.4 (Thermo). Data were searched for peptides with tryptic specificity with a maximum of two missed cleavages. The precursor mass tolerance was set at 10 ppm and the fragment mass tolerance was set at 0.02 Da. Search parameters included carbamidomethylation at cysteine (+57.021 Da) as a constant modification and the following dynamic modifications: oxidation at Met (+15.995 Da), acetylation at protein N-termini (+42.011 Da), Met loss at protein N-termini (−131.040 Da), Met loss+acetylation at protein N-termini (−89.030 Da). The Percolator node of Proteome Discoverer was used for PSM validation at a false discovery rate of 1%. Final results for both LC-MS/MS analyses are given in Supplementary Table S5.

### Mapping ES mass spectrometry hits onto our gene predictions

Our protein database for analyzing ES data included both our new v2.1 *A. ceylanicum* predictions and any predictions by Coghlan *et al*. that differed by even one amino acid residue from ours. We selected ES protein hits in Supplementary Table S5 whose experimental combined q-value was ≤0.01; for each ES sample, we extracted their gene names as separate v2.1 and Coghlan genes, yielding four ES gene lists in all.

Because gene prediction in complex eukaryotes remains challenging (Hatje *et al*. 2019) and because the Coghlan predictions were made on a different strain of *A. ceylanicum* (Indian strain, US National Parasite Collection Number 102954) than our laboratory strain used for genome sequencing and gene prediction (HY135), ES proteins whose mass spectrophotometic data fit a Coghlan prediction exclusively might do so only through slight differences in predicted gene structures or through strain-specific amino acid polymorphisms. To identify such cases where a Coghlan gene was equivalent to one of our v2.1 genes, we mapped the protein-coding exons (CDSes) of Coghlan genes onto our v2.1 gene set, as follows. We downloaded Coghlan gene annotations in GFF3 format and their corresponding *A. ceylanicum* genome assembly from ParaSite (Supplementary Table S3). We mapped the coordinates of Coghlan gene annotations onto our *A. ceylanicum* genome assembly sequence (i.e., lifted them over) with Liftoff 1.6.3 (Shumate and Salzberg 2020) using the arguments ‘*-copies -polish -cds*’. We extracted CDS coordinate lines from the resulting liftover GFF3 file with Perl, and did likewise for the CDS coordinates from the GFF3 of our v2.1 gene prediction set. We converted Coghlan and v2.1 CDS coordinate files from GFF3 to BED format with *gff2bed* in BEDOPS 2.4.41 (Neph *et al*. 2012). We identified overlapping Coghlan and v2.1 CDSes with *intersect* in BEDtools 2.30.0 (Quinlan and Hall 2010), using the argument ‘*-loj*’. We extracted columns 10 and 20 from the resulting intersection file with *cut* (https://www.gnu.org/software/coreutils/cut) using the argument ‘*-f 10,20*’, and processed the results with *washu.cds_loj_umass.cds_genemap.pl* using the positional arguments ‘*[CDS-to-gene table] [BEDtools intersect columns 10 and 20 table]*’; the CDS-to-gene map file was extracted from the FASTA headers of Coghlan and v2.1 proteomes via Perl.

We then used *annot_es_v01.pl* with our Coghlan-to-v2.1 gene map and our four ES gene lists to produce a unified list of ES-encoding v2.1 genes.

### Mapping ES genes of Uzoechi et al. onto our gene predictions

Uzoechi *et al*. have recently identified ES-encoding genes in *A. ceylanicum*, using their own reprediction of our original 2015 gene set (Uzoechi *et al*. 2023). To compare their results to ours, we first mapped their results from ES transcripts to ES genes with *map_tx_to_genes.pl*. We then mapped their predicted gene set onto ours using methods similar to those for the Coghlan gene set. Because their repredictions were performed on our *A. ceylanicum* HY135 genome sequence rather than their *A. ceylanicum* Indian genome (as Coghlan predictions had been), we did not need to use Liftoff to map coordinates from one genome to the other. Subsequent mapping steps (CDS extraction with Perl; GFF to BED conversion with BEDOPS *gff2bed*; overlap determination with BEDtools *intersect*; extraction and summarizing of ES gene results with *annot_es_v02.pl*) were as above.

### Statistical significance of overlapping gene sets

To identify significant overlaps between sets of *A. ceylanicum* genes, we used the Perl script *motif_group_fisher.pl*, which computed p-values from two-tailed Fisher tests with the Perl module *Text::NSP::Measures::2D::Fisher::twotailed*; for multiple hypothesis testing (e.g., comparisons to sets of protein motifs) this Perl script also computed q-values via the *qvalue* program of the MEME 5.4.1 software suite (Bailey *et al*. 2009).

## Results

### Improved A. ceylanicum gene predictions

Analyzing gene function depends crucially on the quality of gene predictions (Hatje *et al*. 2019; GuigÓ 2023). Our published gene set for *A. ceylanicum* (“version 1.0” or “v1.0“; Table 1) used the best resources we had at the time (Schwarz *et al*. 2015), but both software and RNA-seq/protein data for gene prediction have improved since 2015. We thus repredicted *A. ceylanicum* protein-coding genes with BRAKER2 (BrŮna *et al*. 2021), whose predictions we guided with both greatly expanded RNA-seq data sets across the *A. ceylanicum* life cycle (Schwarz *et al*. 2015; Wei *et al*. 2016; Bernot *et al*. 2020) and protein sequences from related nematodes (Doyle *et al*. 2020; Logan *et al*. 2020; Davis *et al*. 2022). We assessed our gene predictions and compared them to others with BUSCO (Waterhouse *et al*. 2018), checking for the presence of 3,131 highly conserved single-copy index genes in nematodes and defining completeness by the percentage of index genes found in a given gene set (Table 1). This gave us an initial repredicted gene set (“version 2.0” or “v2.0”) that was more complete and less fragmented than previous *A. ceylanicum* gene predictions by either us or others. Whereas our v1.0 predictions had detected 87.8% of BUSCO index genes, our v2.0 predictions detected 95.1%. The v2.0 set also had fewer protein-coding genes than v1.0 (19,419 versus 36,687) encoding longer proteins (medians of 300 versus 199 residues), implying that gene predictions in v2.0 were less fragmented than in v1.0. However, we observed that v1.0 included genes that encoded ES proteins (see below) yet were absent from the v2.0 gene set. To detect as many valid ES genes as possible, we identified all v1.0 genes whose exons were completely nonoverlapping with exons of v2.0 genes, and combined them with v2.0 to make a hybrid v2.1 set of 33,190 protein-coding genes. This v2.1 set showed a small but detectable increase in BUSCO completeness over v2.0 (95.1% to 95.3%). Recently, Uzoechi *et al*. have also repredicted *A. ceylanicum* genes using methods similar to ours (Uzoechi *et al*. 2023); their new gene set is also substantially better than earlier predictions (94.5% completeness), though less complete than our v2.0 or v2.1 gene sets (Table 1). Having predicted the v2.1 gene set, we annotated its predicted protein products with predicted N-terminal signal sequences (KÄll *et al*. 2004), conserved protein domains (Mistry *et al*. 2021; Paysan-Lafosse *et al*. 2023), orthologies to genes in related nematode species such as *N. americanus* (Emms and Kelly 2019), and Gene Ontology (GO) terms describing biological and molecular functions (Ashburner *et al*. 2000; Carbon *et al*. 2021); annotations are listed in Supplementary Table S6.

**Table 1.**
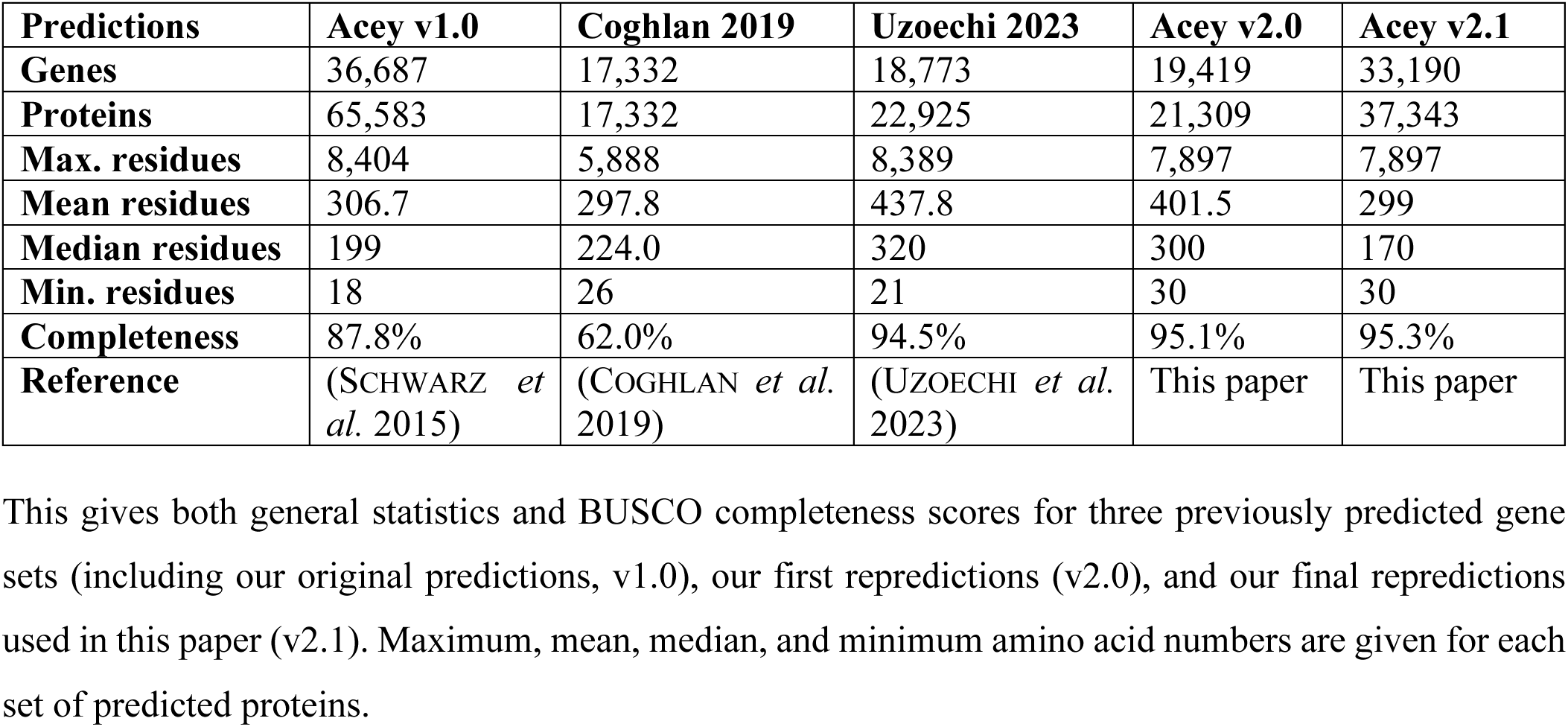
Protein-coding gene predictions for *A. ceylanicum*.

### Identifying ES proteins

To collect ES proteins from adult *A. ceylanicum* hookworms, we isolated hookworms from hamster hosts 20 days after infection in two independent experiments, incubated the hookworms in protein-free culture medium for three days, concentrated and purified their supernatants, subjected their proteins to mass spectrometry, and mapped the resulting protein spectra to a nonredundant set of protein sequences. By this means, we identified 565 genes encoding ES proteins observed in at least one ES collection (1.7% of all genes), and 350 genes encoding ES proteins in both independent collections (1.05% of all genes; Table 2; Supplementary Table S6). Uzoechi *et al*. have also identified genes encoding ES proteins from male and female *A. ceylanicum* (Uzoechi *et al*. 2023); mapping their ES genes onto our v2.1 gene set, we find 860 ES genes from their analysis (2.6% of all genes), of which 430 are identical to ES genes from ours (1.3% of all genes; 76% of our ES genes). This overlap is 29-fold greater than chance (two-tailed Fisher test, p-value = 0), and shows high reproduciblity of ES genes in *A. ceylanicum*. Conversely, each ES gene set has unique members; they collectively have 995 genes (Table 2).

**Table 2.** Numbers of genes encoding ES proteins of *A. ceylanicum*.

| Category | Genes |
| --- | --- |
| Total v2.1 gene set | 33,190 |
| Our ES, at least one collection ($\geq 1$) | 565 |
| Our ES, both collections (2) | 350 |
| Uzoechi ES, either sex | 860 |
| Uzoechi ES, both sexes | 608 |
| Uzoechi ES, male only | 135 |
| Uzoechi ES, female only | 117 |
| Our ES ($\geq 1$) and U. (either) | 430 |
| Our ES (2) and U. (either) | 308 |
| Our ES ($\geq 1$), no U. ES | 135 |
| Our ES (2), no U. ES | 42 |
| Our ES ($\geq 1$), no U. gene | 14 |
| Our ES (2), no U. gene | 7 |
| Either our ES ($\geq 1$) or U. (either) | 995 |
| Either our ES (2) or U. (either) | 902 |

Out of our 565 ES genes, we consider two subsets for detailed analysis here: either categories of ES genes for which we can show statistically significantly enhanced overlaps with other gene categories of interest, or individual instances of ES genes for which we have found specific published information in the literature which indicates that an individual ES gene may have a biologically informative function or effect.

### A. ceylanicum ES genes encode possible host-parasite interaction proteins

Two-thirds of our ES genes (380) were predicted to encode classically secreted proteins with N-terminal signal sequences, 4.6-fold above genome-wide background (q = 4.5•10^−181^); the remaining one-third of ES genes (185) encode proteins that are presumably non-classically secreted (Table 3; Supplementary Table S7) (Dimou and Nickel 2018). This mix of predominant but not universal N-terminal signals also exists in ES proteins from *A. caninum*, *N. americanus*, and *H. contortus* (Supplementary Tables S8-S10) (Morante *et al*. 2017; Wang *et al*. 2019; Logan *et al*. 2020), and may reflect nonclassical secretion of ES proteins through extracellular vesicles (Balmer and Faso 2021; Wang *et al*. 2024).

**Table 3.** Protein motifs overrepresented in *A. ceylanicum* ES genes.

| **Motif** | **Motif genes** | **Overlap** | **Enrichment** | **q-value** |
| --- | --- | --- | --- | --- |
| N-terminal signal sequence (SigP) | 4,825 | 380 | 4.63 | 4.51e-181 |
| Cysteine-rich secretory protein (CAP) [PF00188.29] | 418 | 140 | 19.67 | 5.64e-140 |
| ASP-related (ASPR) [PF17641.5] | 200 | 51 | 14.98 | 1.88e-41 |
| Papain family cysteine protease [PF00112.26] | 67 | 24 | 21.04 | 1.60e-22 |
| Trypsin inhibitor-like cysteine-rich domain (TIL) [PF01826.20] | 71 | 19 | 15.72 | 7.83e-15 |
| Astacin (Peptidase family M12A) [PF01400.27] | 122 | 22 | 10.59 | 1.41e-13 |
| Kunitz/bovine pancreatic trypsin inhibitor domain [PF00014.26] | 111 | 19 | 10.06 | 3.74e-11 |
| Transthyretin-like family (TTL, e.g., TTR-52) [PF01060.26] | 87 | 16 | 10.8 | 1.01e-09 |
| Potassium channel inhibitor ShK domain-like [PF01549.27] | 111 | 16 | 8.47 | 4.18e-08 |
| Tissue inhibitor of metalloproteinase (TIMP) [PF00965.20] | 31 | 8 | 15.16 | 0.0000169 |
| Secreted clade V proteins (SCVP) [PF17619.5] | 59 | 10 | 9.96 | 0.000023 |
| Glutathione S-transferase, C-terminal domain [PF00043.28] | 22 | 5 | 13.35 | 0.0109 |
| Peptidase family M13/neprilysin metallopeptidase [PF01431.24] | 39 | 6 | 9.04 | 0.0163 |
| Aspartyl protease [PF00026.26] | 57 | 7 | 7.21 | 0.0163 |
| Lectin C-type domain [PF00059.24] | 102 | 9 | 5.18 | 0.0191 |
| Serine carboxypeptidase S28 [PF05577.15] | 14 | 4 | 16.78 | 0.0196 |
| Peptidase family M13/neprilysin metallopeptidase, N-terminal domain [PF05649.16] | 42 | 6 | 8.39 | 0.0196 |

Compared to the whole *A. ceylanicum* proteome, ES genes disproportionately encoded several protein families with plausible roles in host-parasite interactions (Table 3; Supplementary Table S7) (Abuzeid *et al*. 2020). These included five types of proteases (aspartyl, astacin, cysteine, serine, and metallopeptidase) that may digest host tissues and blood (Williamson *et al*. 2006; Ranjit *et al*. 2009; Knox 2012; Yang *et al*. 2015; Caffrey *et al*. 2018), along with three types of protease inhibitors (Kunitz, TIL, and TIMP) that may protect *A. ceylanicum* from native or host proteases; both proteases and protease inhibitors may also counteract host immune responses (Chu *et al*. 2004; Knox 2007). Five ES genes encoded glutathione S-transferase, an enzyme thought to detoxify free heme generated during hemoglobin proteolysis and other toxins (Matouskova *et al*. 2016; Abuzeid *et al*. 2020). Sixteen ES genes encoded *Stichodactyla helianthus* toxin (ShK)-related proteins, which might suppress host T cells (Chhabra *et al*. 2014; McNeilly *et al*. 2017); and nine ES genes encoded C-type lectin proteins, which might mimic mammalian immune proteins (enabling immunosuppression) or dissociate cells (enabling tissue invasion) (Loukas *et al*. 2000; Harcus *et al*. 2009; Yoshida *et al*. 2012; Schwarz *et al*. 2015; Wang *et al*. 2023).

One quarter of ES genes (140) encoded secreted venom-like allergen proteins with CAP or SCP/TAPS domains (Gibbs *et al*. 2008; Cantacessi *et al*. 2009); in hookworms, these are called *Ancylostoma* secreted proteins, activation-associated secreted proteins, or ASPs (Hawdon *et al*. 1996; Yatsuda *et al*. 2002; Wilbers *et al*. 2018). One ASP, neutrophil inhibitory factor in *A. caninum* (*Acan*-NIF), has been experimentally shown to block neutrophil adhesion (Moyle *et al*. 1994; Lo *et al*. 1999); another ASP, hookworm platelet inhibitor in *A. caninum* (*Acan*-HPI), has been shown to prevent blood clotting (Del Valle *et al*. 2003). We observed ES genes encoding orthologs of both these ASPs (*Acey*-NIF-B and *Acey*-HPI) which may themselves be immunosuppressant and antithrombotic proteins (Supplementary Table S6). However, this leaves the other 138 ES ASP genes with only conjectural functions.

ES genes also encode three other multigene families of secreted proteins that are suspected to be involved in infection, although their molecular roles remain largely uncharacterized: ASP-related (ASPR), transthyretin-like (TTL), and secreted clade V proteins (SCVP). ASPRs were first observed in *A. ceylanicum* as a divergent subfamily of ASPs transcriptionally upregulated immediately after host infection (Schwarz *et al*. 2015), and were later observed in ES proteins from *N. americanus* (Logan *et al*. 2020). TTL proteins were previously observed as a major component of ES proteins from *H. contortus* (Wang *et al*. 2019) and as multigene families in several parasitic nematodes (Hunt *et al*. 2017; Tritten *et al*. 2021; Stevens *et al*. 2023); one TTL protein of *H. contortus* (*Hc*TTR) antagonizes goat IL-4 *in vitro*, and thus may be immunosuppressive *in vivo* (Tian *et al*. 2019). SCVPs were observed in *A. ceylanicum* as a novel family of secreted proteins upregulated in young hookworm adults that is conserved in *C. elegans* and expanded in hookworms and other strongylids (Schwarz *et al*. 2015).

Other ES genes were potentially relevant to infection despite not being from overrepresented families (Table 4). One encodes a macin homolog that may protect hookworms from being infected by bacteria, and thus enable their long-term residence in the human gut (Jung *et al*. 2012; Irvine *et al*. 2023). Three ES homologs of *A. caninum* anticoagulant protein (Stassens *et al*. 1996) could act with *Acey*-HPI to prevent clotting during blood feeding. Seven other ES genes encode possible immunosuppressants: two ES apyrases and one ES adenosine deaminase could suppress both blood clotting and inflammation by hydrolyzing extracellular ATP (Gounaris and Selkirk 2005; Guiguet *et al*. 2016; Pala *et al*. 2023); phospholipase A2, phospolipase D, and fatty acid- and retinol-binding (FAR) ES proteins could suppress lipid signals in immune responses (Parks *et al*. 2021; Parks *et al*. 2023); the ES macrophage migration inhibitory factor *Acey*-MIF-1, previously observed to bind the human MIF receptor CD74 and promote monocyte migration, could alter cellular immune responses (Cho *et al*. 2007; Sparkes *et al*. 2017); and an ES deoxyribonuclease II could hydrolyze extracellular DNA nets secreted by neutrophils to trap and kill large pathogens (Bouchery *et al*. 2020).

**Table 4.** Individual *A. ceylanicum* ES genes of interest.

| **Predicted product** | **ES protein-encoding gene** | **Possible importance** |
| --- | --- | --- |
| *A. caninum* anticoagulant protein (AcAP) homolog | Acey_s0112.v2.g13097 | Anticoagulant |
| *A. caninum* anticoagulant protein (AcAP) homolog | Acey_s0112.v2.g13104 | Anticoagulant |
| *A. caninum* anticoagulant protein (AcAP) homolog | Acey_s0112.v2.g13105 | Anticoagulant |
| Adenosine deaminase | Acey_s0243.v2.g530 | Anti-coagulation/-inflammation |
| Apyrase | Acey_s0038.v2.g9677 | Anti-coagulation/-inflammation; vaccine subunit |
| Apyrase | Acey_s0038.v2.g9678 | Anti-coagulation/-inflammation; vaccine subunit |
| Calreticulin | Acey_s0340.v2.g13758 | Vaccine subunit |
| Cysteine protease | Acey-CP-1 | Digestion; vaccine subunit |
| Deoxyribonuclease II | Acey_s0473.v2.g18261 | Hydrolyzing neutrophil DNA nets |
| Enolase | Acey_s0389.v2.g8512 | Vaccine subunit |
| Fatty acid- and retinol-binding (FAR) protein | Acey_s0005.v2.g14583 | Lipid immunosuppression; vaccine subunit |
| Fatty acid- and retinol-binding (FAR) protein | Acey_s0064.v2.g6025 | Lipid immunosuppression; vaccine subunit |
| Fructose-1,6-bisphosphate aldolase | Acey_s0324.v2.g12031 | Vaccine subunit |
| Glutathione S-transferase | Acey-GST-1 | Detoxification; vaccine subunit |
| Hypodermal Ac-16 antigen homolog | Acey_s0009.v2.g6653 | Vaccine subunit |
| M13 peptidase | Acey-MEP-6 | Digestion; vaccine subunit |
| M13 peptidase | Acey-MEP-7 | Digestion; vaccine subunit |
| Macin | Acey_s0022.v2.g18553 | Antimicrobial peptide |
| Macrophage migration inhibitory factor | Acey-MIF-1 | Suppressing cellular immunity |
| Phospholipase A2 | Acey_s0002.v2.g4275 | Lipid immunosuppression |
| Phospholipase D | Acey_s0097.v2.g996 | Lipid immunosuppression |

Because ES proteins are exposed to the human host’s immune system and are likely to promote infection, ES proteins are promising targets for an anti-hookworm vaccine. Twelve ES genes encoded proteins that have already been shown to give partial protection as vaccines in laboratory mammals, or to be homologous to such proteins (Table 4): the cysteine protease *Acey*-CP-1 (Noon *et al*. 2019); the M13 peptidases *Acey*-MEP-6 and *Acey*-MEP-7 (Wisniewski *et al*. 2013; Wisniewski *et al*. 2016); the glutathione S-transferase *Acey*-GST-1, homologous to *A. caninum*’s *Acan*-GST-1 (Xiao *et al*. 2008) and *N. americanus’ Na*-GST-1 (Zhan *et al*. 2010); a homolog of the immunodominant hypodermal *Ac*-16 antigen of *A. caninum* (Fujiwara *et al*. 2007); two homologs of apyrases in *Teledorsagia circumcincta* and *H. bakeri* (Nisbet *et al*. 2019; Berkachy *et al*. 2021); two homologs of fatty acid- and retinol-binding proteins in *A. ceylanicum* (Fairfax *et al*. 2009) and *Onchocerca volvulus* (Jenkins *et al*. 1996); one homolog of *Brugia malayi* calreticulin (Yadav *et al*. 2021); one homolog of *Onchocerca volvulus* fructose-1,6-bisphosphate aldolase (McCarthy *et al*. 2002); and one homolog to *A. suum* enolase (Chen *et al*. 2012). In addition to these strict orthologs, three instances of astacin, ASP, and TTL proteins (all overrepresented among ES proteins) have given partial protection as vaccines: *Ac*-MTP-1 of *A. caninum*, *Acan*-ASP-2 of *A. caninum*, and *Hc*TTR of *H. contortus* (Zhan *et al*. 2002; Bethony *et al*. 2005; Tian *et al*. 2020). These results show the vaccine potential of ES proteins, with hundreds of other ES proteins remaining untested as vaccine subunits.

Over half of *A. ceylanicum* ES genes are functionally conserved in related blood-feeding parasitic nematodes: they do not merely have homologs, but ES gene homologs (Table 5; Supplementary Tables S6 and S11). Out of 565 *A. ceylanicum* ES genes, 269 (47.6%) had ES gene homologs in the closely related hookworm *A. caninum* (Morante *et al*. 2017), 21-fold more than expected by chance (p = 6.82•10^−301^). For the more distantly related hookworm *N. americanus* (Logan *et al*. 2020), 141/565 ES genes (25.0%) had ES gene homologs (18-fold over chance; p = 1.78•10^−137^); for the still more distantly related blood-feeding *H. contortus* (Wang *et al*. 2019), 180/565 ES genes (31.9%) had ES gene homologs (8.9-fold over chance, p = 4.10•10^−120^). Considering all three species at once, 350/565 ES genes (61.9%) had some ES homolog (12-fold over chance, p = 0). Similarly high conservation was observed for the 860 ES genes described by Uzoechi *et al*. (Supplementary Table S11) (Uzoechi *et al*. 2023). These homologies of *A. ceylanicum* ES genes in other species are predominantly to ES genes themselves (i.e., genes experimentally shown to encode ES proteins in these other species). When ES genes in other species are set aside, the remaining homologies of *A. ceylanicum* ES genes fall to, or even below, background frequencies. For *A. ceylanicum* ES genes without known ES gene homologs, homology to *A. caninum* dropped to 1.6-fold over chance (p = 1.31•10^−29^); in *N. americanus*, to 1.1-fold below chance (p = 0.22); and in *H. contortus*, to 2.4-fold below chance (p = 2.05•10^−30^). Thus, the *A. ceylanicum* ES gene set predominantly encodes ES proteins not only in *A. ceylanicum* itself but also in related parasitic nematodes.

### Identification of intestinal and immunoregulated genes

Because the hookworm intestine is an important source of ES proteins, we set out to characterize intestine- and non-intestine-biased expression of hookworm genes. We infected hamsters with *A. ceylanicum* hookworms, allowed infections to proceed for either 12 or 19 days, collected hookworms from hamster small intestines, extracted hookworm RNAs, and performed RNA-seq with 36.9 to 55.8 million reads per biological replicate (Supplementary Table S1). Hookworms at the 12-day infection stage were young adults that were too small for us to dissect further, so RNA-seq was done on whole 12-day hookworms. By 19 days, hookworms had grown into mature adults large enough to dissect, so we separated them into intestinal and non-intestinal tissues before RNA-seq. In addition, we hypothesized that hookworms would respond to the mammalian host’s immune system by increasing their expression of proteins that counteract the mammalian immune system. To detect hookworm genes regulated by the state of the host’s immune system, 12-day and 19-day infections were performed both with normal hamster hosts (nonDEX) and with hamsters whose immune systems had been suppressed by dexamethasone (DEX). We analyzed levels and changes of gene expression both for our RNA-seq data and for published *A. ceylanicum* RNA-seq data from adult male intestine, male adults, and female adults (Table 6; Figure 2; Supplementary Table S6). We defined a gene as significantly changing its activity between two biological conditions (e.g., intestinal versus non-intestinal 19-day tissues, or intestinal nonDEX versus intestinal DEX) if the gene changed its expression by at least 2-fold (log2FC ≤ −1 or ≥ 1) with a false discovery rate (p-value corrected for multiple hypothesis testing) of no more than 0.01 (FDR ≤ 0.01).

**Table 5.**
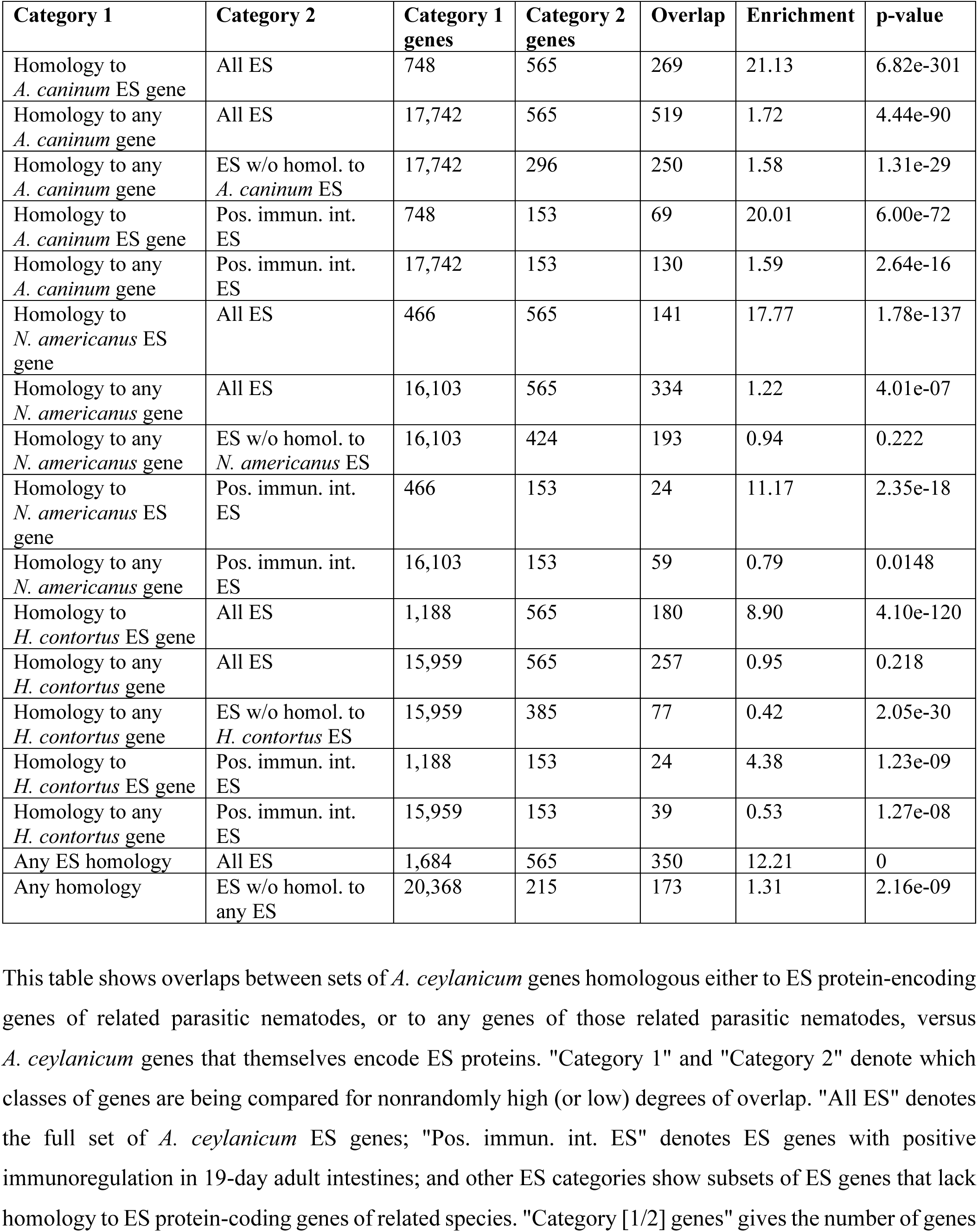
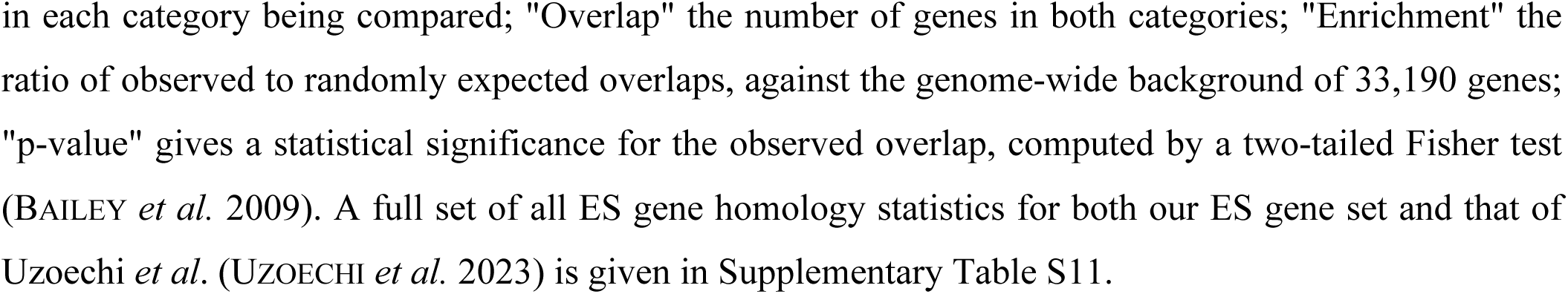
Homologies of ES genes of *A. ceylanicum* to those of related parasitic nematodes.

| Category 1 | Category 2 | Category 1 genes | Category 2 genes | Overlap | Enrichment | p-value |
| --- | --- | --- | --- | --- | --- | --- |
| Homology to *A. caninum* ES gene | All ES | 748 | 565 | 269 | 21.13 | 6.82e-301 |
| Homology to any *A. caninum* gene | All ES | 17,742 | 565 | 519 | 1.72 | 4.44e-90 |
| Homology to any *A. caninum* gene | ES w/o homol. to *A. caninum* ES | 17,742 | 296 | 250 | 1.58 | 1.31e-29 |
| Homology to *A. caninum* ES gene | Pos. immun. int. ES | 748 | 153 | 69 | 20.01 | 6.00e-72 |
| Homology to any *A. caninum* gene | Pos. immun. int. ES | 17,742 | 153 | 130 | 1.59 | 2.64e-16 |
| Homology to *N. americanus* ES gene | All ES | 466 | 565 | 141 | 17.77 | 1.78e-137 |
| Homology to any *N. americanus* gene | All ES | 16,103 | 565 | 334 | 1.22 | 4.01e-07 |
| Homology to any *N. americanus* gene | ES w/o homol. to *N. americanus* ES | 16,103 | 424 | 193 | 0.94 | 0.222 |
| Homology to *N. americanus* ES gene | Pos. immun. int. ES | 466 | 153 | 24 | 11.17 | 2.35e-18 |
| Homology to any *N. americanus* gene | Pos. immun. int. ES | 16,103 | 153 | 59 | 0.79 | 0.0148 |
| Homology to *H. contortus* ES gene | All ES | 1,188 | 565 | 180 | 8.90 | 4.10e-120 |
| Homology to any *H. contortus* gene | All ES | 15,959 | 565 | 257 | 0.95 | 0.218 |
| Homology to any *H. contortus* gene | ES w/o homol. to *H. contortus* ES | 15,959 | 385 | 77 | 0.42 | 2.05e-30 |
| Homology to *H. contortus* ES gene | Pos. immun. int. ES | 1,188 | 153 | 24 | 4.38 | 1.23e-09 |
| Homology to any *H. contortus* gene | Pos. immun. int. ES | 15,959 | 153 | 39 | 0.53 | 1.27e-08 |
| Any ES homology | All ES | 1,684 | 565 | 350 | 12.21 | 0 |
| Any homology | ES w/o homol. to any ES | 20,368 | 215 | 173 | 1.31 | 2.16e-09 |

This table shows overlaps between sets of *A. ceylanicum* genes homologous either to ES protein-encoding genes of related parasitic nematodes, or to any genes of those related parasitic nematodes, versus *A. ceylanicum* genes that themselves encode ES proteins. "Category 1" and "Category 2" denote which classes of genes are being compared for nonrandomly high (or low) degrees of overlap. "All ES" denotes the full set of *A. ceylanicum* ES genes; "Pos. immun. int. ES" denotes ES genes with positive immunoregulation in 19-day adult intestines; and other ES categories show subsets of ES genes that lack homology to ES protein-coding genes of related species. "Category [1/2] genes" gives the number of genes

**Table 6.** Numbers of *A. ceylanicum* genes showing differential RNA expression (significant changes of gene expression between biological conditions).

| Category | Genes |
| --- | --- |
| Total v2.1 gene set | 33,190 |
| Intestine-biased (19-day) | 1,670 |
| Non-intestine-biased (19-day) | 1,196 |
| Intestine-biased (prev. pub. data) | 635 |
| Non-intestine-biased (prev. pub. data) | 954 |
| 19-day positively immunoregulated intestinal | 1,951 |
| 19-day negatively immunoregulated intestinal | 137 |
| 19-day positively non-intestinal immunoreg. | 26 |
| 19-day negatively non-intestinal immunoreg. | 4 |
| Any 19-day immunoregulated | 2,112 |
| 12-day positively immunoregulated | 0 |
| 12-day negatively immunoregulated | 2 |
| Male-biased (prev. pub. data) | 3,135 |
| Female-biased (prev. pub. data) | 1,547 |
| Any change of gene expression above | 8,130 |

**Figure 2.**
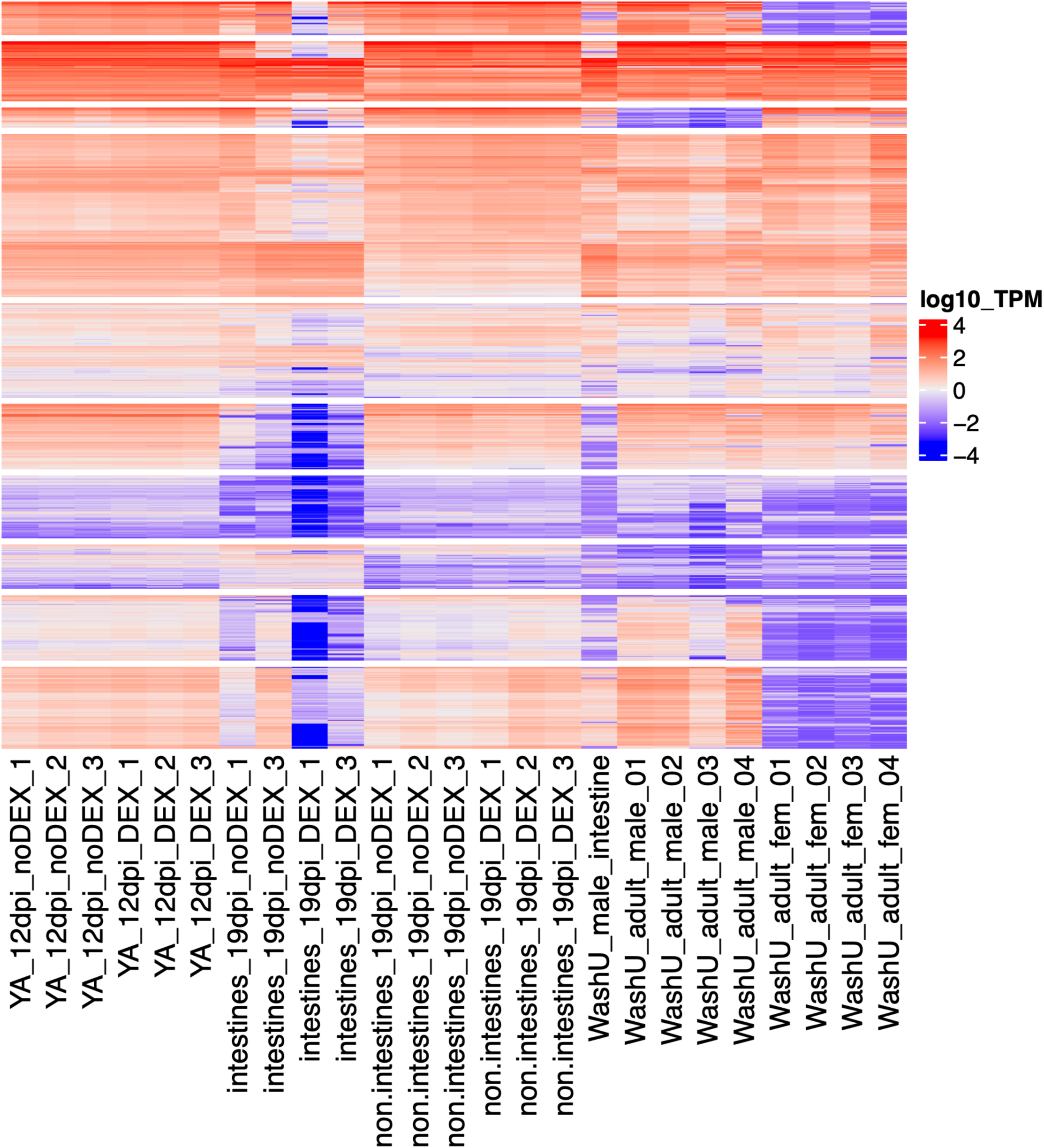
Gene expression in *A. ceylanicum*. Gene activity is shown for 8,130 *A. ceylanicum* genes with significant differences in expression between biological conditions, with biological replicates used for differential gene expression analysis on the x-axis and individual genes on the y-axis. Expression levels are in TPM (log10). Genes sharing similar patterns of expression are split into 10 clusters. Biological replicates are: young 12-day hookworm adults from normal hosts (YA_12dpi_noDEX); young 12-day adults from immunosuppressed hosts (YA_12dpi_DEX); dissected 19-day intestines from mature hookworm adults in normal hosts (intestines_19dpi_noDEX); dissected 19-day intestines in immunosuppressed hosts (intestines_19dpi_DEX); dissected 19-day intestines in normal hosts (non.intestines_19dpi_noDEX); dissected 19-day non-intestinal tissues in immunosuppressed hosts (non.intestines_19dpi_DEX); previously published adult male intestine (WashU_male_intestine); published adult males (WashU_adult_male); and published adult females (WashU_adult_fem). Additional heatmaps of gene expression for all 33,190 genes and all replicates are given in Supplemental Figure 1.

By these criteria, in 19-day adults we observed 1,670 genes with intestine-biased expression and 1,196 with non-intestine-biased expression, the remaining 30,324 genes being intermediate (Table 6; Figure 2). Of the intestine-biased genes, 723 (43.3%) were homologous to *H. contortus* genes with intestine-biased expression (4.4-fold above background; p = 8.18•10^−306^; Supplemental Table S15). For comparison, when we reanalyzed published RNA-seq data for male intestine versus whole male adults (Wei *et al*. 2016; Bernot *et al*. 2020) as part of the same computation, we identified 635 genes with intestine-biased expression, of which only 180 were also found in our intestine-biased gene set (Supplementary Table S13). By chance alone, one would expect to observe 32 genes shared by both intestinal sets; the observed overlap was 5.6-fold above background (p = 9.93•10^−85^) but limited (only 28% of the 635 genes were shared). The observation of over 700 genes with intestine-biased expression conserved between strongylids supports our larger gene set. The disagreement of these intestine-biased gene sets could have one or more causes: different protocols for dissection and RNA extraction; different background tissues for comparison to intestinal RNA-seq (non-intestinal here, versus whole adults in the previous data); different sexes (mixed-sex here, versus males previously); and different strains of *A. ceylanicum* (HY135 here, versus Indian previously), which could cause different rates of RNA-seq read mapping onto our HY135 reference genome (Supplementary Table S12) (Degner *et al*. 2009; Rezansoff *et al*. 2019; Bell *et al*. 2023).

When comparing *A. ceylanicum* in normal (nonDEX) versus immunosuppressed (DEX) hosts, we detected substantial changes of gene expression in 19-day hookworm intestinal tissues, but almost no changes of gene expression either in 12-day young adults or in 19-day non-intestinal tissues. We observed 1,951 genes upregulated in 19-day nonDEX intestine versus 19-day DEX intestine (5.9% of all genes), along with 137 downregulated genes (0.41%). In contrast, we observed only 32 genes positively or negatively immunoregulated either in 12-day young adults or in 19-day non-intestinal tissues (Table 6; Figure 2). One explanation for this pattern is that only 19-day intestinal tissues are directly exposed to the host’s immune system through blood feeding. Although 12-day young adult hookworms inhabit the small intestine, they have only just started feeding on blood; up to that point, they instead probably eat mucosal cells (Bansemir and Sukhdeo 1994; Bansemir and Sukhdeo 2001). By 19 days, hookworm adults have been feeding on blood for up to a week; however, most of their bodily contact with this blood (and, thus, with the host’s immune system) is through the lumen of their intestine (although cephalic or pharyngeal glands may interact with their host as well).

### Intestine-biased genes encode likely components of food digestion, detoxification, and absorption

Whereas ES genes overwhelmingly encoded proteins with N-terminal signal sequences alone, intestine-biased genes were modestly but significantly enriched for several predicted types of secreted or transmembrane proteins (Table 7; Supplementary Table S13). These included secreted proteins, proteins predicted to have one transmembrane anchor sequence after the N-terminal signal peptide, and multi-transmembrane proteins. Intestinal genes disproportionately encoded protein families relevant to digesting, detoxifying, and absorbing food (Table 7): aspartyl, cysteine, aminopeptidase, and metallopeptidase proteases (Williamson *et al*. 2006; Ranjit *et al*. 2009; Knox 2012; Yang *et al*. 2015; Caffrey *et al*. 2018); UDP-glucoronosyl or UDP-glucosyl transferases, ecdysteroid kinase-like enzymes, and ABC transporters (Matouskova *et al*. 2016; Scanlan *et al*. 2022; Raza *et al*. 2023); and major facilitator and sugar transporters (Chen *et al*. 2015; Drew *et al*. 2021). Intestinal genes also disproportionately encoded histones; this raises the question of whether mitosis persists in the adult hookworm intestine, as has been observed or inferred for *A. suum* and *H. bakeri* (Anisimov and Tokmakova 1974; Anisimov and Usheva 1974; Pollo *et al*. 2024). In addition to these well-studied protein families, another notably overrepresented family was Strongylid L4 proteins (SL4Ps), first observed in *A. ceylanicum* as a novel family of non-classically secreted proteins upregulated in fourth-stage (L4) larvae (Schwarz *et al*. 2015). Analyzing previously published *H. contortus* intestinal RNA-seq data (Supplementary Table S4), we observed that SL4P genes were also disproportionately represented among intestine-biased *H. contortus* genes, along with several of the better-characterized protein families implicated in digestion, detoxification, or absorption (e.g., cysteine proteases, UDP-glucoronosyl/glucosyl transferases, and major facilitator transporters; Supplementary Table S14). Moreover, the *C. elegans* genes *numr-1* and *numr-2* encode SL4P proteins that are intestinally expressed (Tvermoes *et al*. 2010). We conclude that SL4Ps have conserved intestinal expression in hookworms and other nematodes, and that (as in *C. elegans*) their function might be to counteract toxic environmental stresses (Wu *et al*. 2019; Hong *et al*. 2024).

**Table 7.** Protein motifs overrepresented in *A. ceylanicum* genes with intestine-biased expression.

| Motif | Motif genes | Overlap | Enrichment | q-value |
| --- | --- | --- | --- | --- |
| Papain family cysteine protease [PF00112.26] | 67 | 48 | 14.24 | 1.08e-43 |
| Eukaryotic aspartyl protease [PF00026.26] | 57 | 30 | 10.46 | 7.40e-21 |
| Xylanase inhibitor N-terminal [PF14543.9] | 43 | 22 | 10.17 | 1.37e-14 |
| Uncharacterized oxidoreductase *dhs-27* (DUF1679) [PF07914.14] | 19 | 15 | 15.69 | 1.14e-13 |
| Ecdysteroid kinase-like family [PF02958.23] | 18 | 14 | 15.46 | 1.44e-12 |
| UDP-glucuronosyl and UDP-glucosyl transferase [PF00201.21] | 48 | 21 | 8.70 | 2.18e-12 |
| Major facilitator superfamily [PF07690.19] | 149 | 32 | 4.27 | 1.63e-09 |
| Phosphotransferase enzyme family [PF01636.26] | 34 | 15 | 8.77 | 1.33e-08 |
| Core histone H2A/H2B/H3/H4 [PF00125.27] | 53 | 18 | 6.75 | 2.35e-08 |
| SigP+TM(1x) | 673 | 72 | 2.13 | 7.83e-08 |
| Glycosyltransferase family 28 C-terminal domain [PF04101.19] | 39 | 15 | 7.64 | 1.14e-07 |
| TM(11x) | 111 | 23 | 4.12 | 1.22e-07 |
| Histone-like transcription factor (CBF/NF-Y) and archaeal histone [PF00808.26] | 54 | 16 | 5.8 | 2.21e-06 |
| SigP | 4,825 | 318 | 1.31 | 3.02e-06 |
| Sugar (and other) transporter [PF00083.27] | 71 | 18 | 5.04 | 3.56e-06 |
| ABC transporter [PF00005.30] | 73 | 18 | 4.90 | 5.29e-06 |
| TM(12x) | 140 | 23 | 3.27 | 5.34e-06 |
| TM(10x) | 99 | 18 | 3.61 | 1.60e-05 |
| Strongyloid L4 protein (SL4P) [PF17618.5] | 25 | 10 | 7.95 | 5.33e-05 |
| TM(6x) | 237 | 29 | 2.43 | 9.55e-05 |
| C-terminus of histone H2A [PF16211.8] | 18 | 8 | 8.83 | 0.000338 |
| Peptidase family M1/aminopeptidase [PF01433.23] | 24 | 9 | 7.45 | 0.000375 |
| Peptidase family M13/neprilysin metallopeptidase [PF01431.24] | 39 | 11 | 5.61 | 0.000611 |

### Immunoregulated genes encode signal transduction, male-associated, host-parasite, and ES proteins

The 1,951 positively immunoregulated intestinal genes were modestly, but significantly, enriched for encoding secreted proteins, proteins with a single transmembrane anchor, or both: in other words, possible secreted or cell-surface proteins (Table 8). Overrepresented protein families included possible signal transduction components: protein kinases, protein phosphatases, and SH2 scaffold proteins (Taylor and Kornev 2011; Diop *et al*. 2022; Kokot and Kohn 2022). Other overrepresented families included homologs of nematode motile sperm proteins (MSPs) (Smith 2014), along with two other classes of proteins associated with MSPs: MSP fiber 2 proteins (MFP2s) (Grant *et al*. 2005) and DUF236 proteins (RÖdelsperger *et al*. 2021). Although MSPs are indeed hallmark proteins of nematode sperm cells and are usually assumed to be entirely specific to sperm, there are in fact MSPs expressed in somatic cells and required for their mobility. Two different MSP homologs in *Caenorhabditis elegans* have cell and axonal migration RNAi phenotypes in male linker cells and hermaphroditic neurons, with one of them being expressed in linker cells (Schmitz *et al*. 2007; Schwarz *et al*. 2012). Moreover, *msp* genes are expressed in *C. elegans* ADL chemosensory neurons (Ow *et al*. 2024), and are evolutionarily retained even in parthenogenetic nematode species lacking sperm (Heger *et al*. 2010). Thus, nematodes express some *msp* genes somatically, which fits our observation here of *msp*, *mfp2*, and *duf236* gene expression in dissected intestines of adult *A. ceylanicum*. In addition, positively immunoregulated intestinal genes disproportionately encoded protein families with possible roles in host-parasite interactions. These families included: astacin, leishmanolysin, metallopeptidase, and trypsin-like proteases; TIMP protease inhibitors; ShK-related proteins; ASPs and ASPRs; SCVPs; and immunoglobulin domain-containing proteins (Table 8). While most of these families were also enriched among ES genes, leishmanolysins, trypsins, and immunoglobulin domains were uniquely enriched here. Despite being expressed intestinally, only one of these immunoregulated genes had intestine-biased expression; instead, 258 were non-intestine-biased (3.67-fold over chance; p = 5.51•10^−78^; Supplementary Table S15).

**Table 8.**
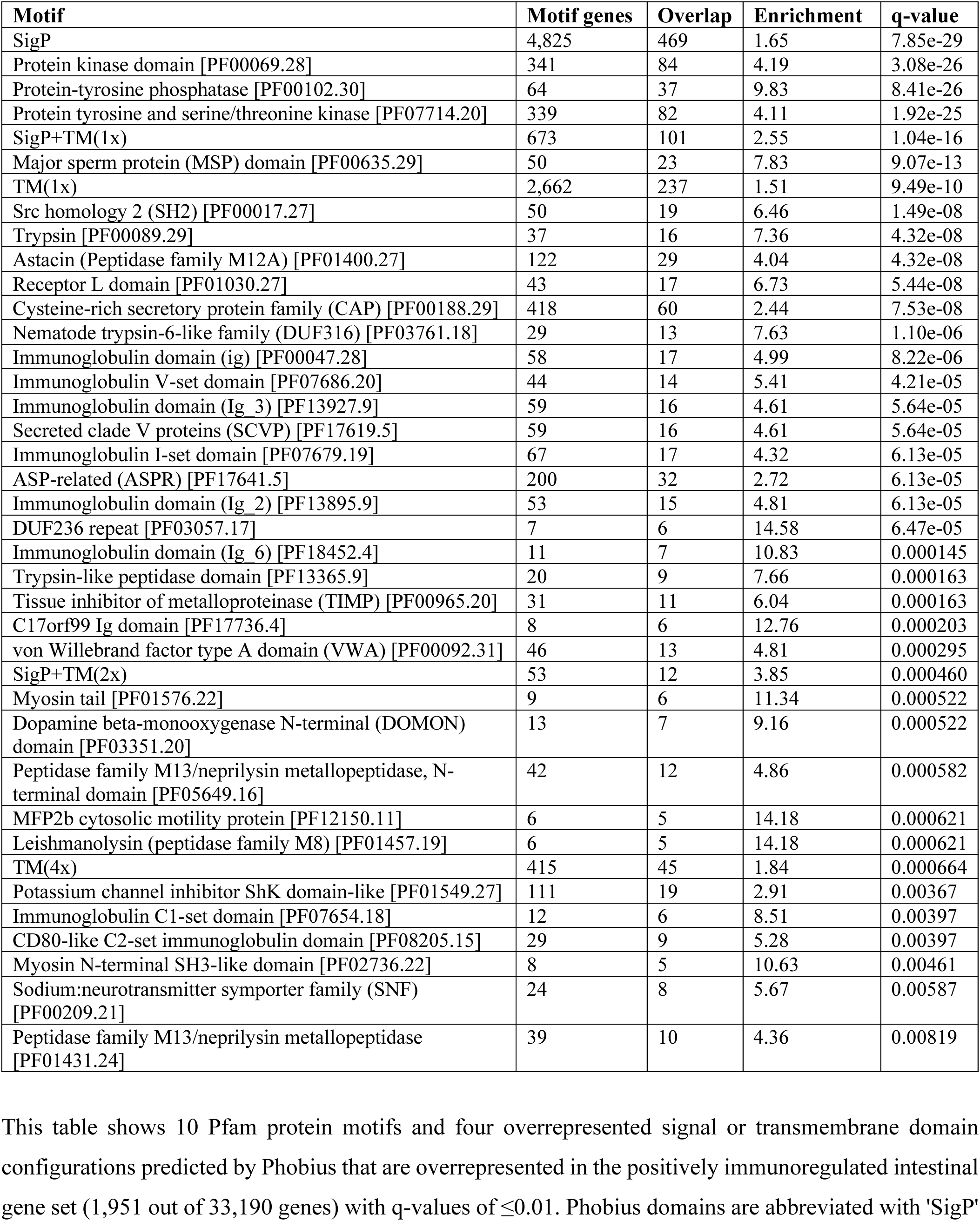
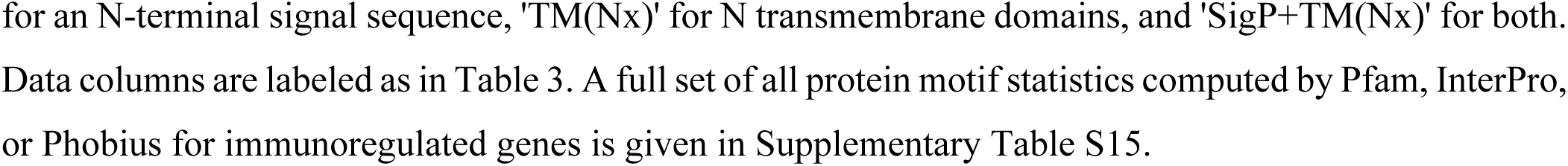
Protein motifs overrepresented in positively immunoregulated intestinal *A. ceylanicum* genes.

| Motif | Motif genes | Overlap | Enrichment | q-value |
| --- | --- | --- | --- | --- |
| SigP | 4,825 | 469 | 1.65 | 7.85e-29 |
| Protein kinase domain [PF00069.28] | 341 | 84 | 4.19 | 3.08e-26 |
| Protein-tyrosine phosphatase [PF00102.30] | 64 | 37 | 9.83 | 8.41e-26 |
| Protein tyrosine and serine/threonine kinase [PF07714.20] | 339 | 82 | 4.11 | 1.92e-25 |
| SigP+TM(1x) | 673 | 101 | 2.55 | 1.04e-16 |
| Major sperm protein (MSP) domain [PF00635.29] | 50 | 23 | 7.83 | 9.07e-13 |
| TM(1x) | 2,662 | 237 | 1.51 | 9.49e-10 |
| Src homology 2 (SH2) [PF00017.27] | 50 | 19 | 6.46 | 1.49e-08 |
| Trypsin [PF00089.29] | 37 | 16 | 7.36 | 4.32e-08 |
| Astacin (Peptidase family M12A) [PF01400.27] | 122 | 29 | 4.04 | 4.32e-08 |
| Receptor L domain [PF01030.27] | 43 | 17 | 6.73 | 5.44e-08 |
| Cysteine-rich secretory protein family (CAP) [PF00188.29] | 418 | 60 | 2.44 | 7.53e-08 |
| Nematode trypsin-6-like family (DUF316) [PF03761.18] | 29 | 13 | 7.63 | 1.10e-06 |
| Immunoglobulin domain (ig) [PF00047.28] | 58 | 17 | 4.99 | 8.22e-06 |
| Immunoglobulin V-set domain [PF07686.20] | 44 | 14 | 5.41 | 4.21e-05 |
| Immunoglobulin domain (Ig 3) [PF13927.9] | 59 | 16 | 4.61 | 5.64e-05 |
| Secreted clade V proteins (SCVP) [PF17619.5] | 59 | 16 | 4.61 | 5.64e-05 |
| Immunoglobulin I-set domain [PF07679.19] | 67 | 17 | 4.32 | 6.13e-05 |
| ASP-related (ASPR) [PF17641.5] | 200 | 32 | 2.72 | 6.13e-05 |
| Immunoglobulin domain (Ig 2) [PF13895.9] | 53 | 15 | 4.81 | 6.13e-05 |
| DUF236 repeat [PF03057.17] | 7 | 6 | 14.58 | 6.47e-05 |
| Immunoglobulin domain (Ig 6) [PF18452.4] | 11 | 7 | 10.83 | 0.000145 |
| Trypsin-like peptidase domain [PF13365.9] | 20 | 9 | 7.66 | 0.000163 |
| Tissue inhibitor of metalloproteinase (TIMP) [PF00965.20] | 31 | 11 | 6.04 | 0.000163 |
| C17orf99 Ig domain [PF17736.4] | 8 | 6 | 12.76 | 0.000203 |
| von Willebrand factor type A domain (VWA) [PF00092.31] | 46 | 13 | 4.81 | 0.000295 |
| SigP+TM(2x) | 53 | 12 | 3.85 | 0.000460 |
| Myosin tail [PF01576.22] | 9 | 6 | 11.34 | 0.000522 |
| Dopamine beta-monooxygenase N-terminal (DOMON) domain [PF03351.20] | 13 | 7 | 9.16 | 0.000522 |
| Peptidase family M13/neprilysin metallopeptidase, N-terminal domain [PF05649.16] | 42 | 12 | 4.86 | 0.000582 |
| MFP2b cytosolic motility protein [PF12150.11] | 6 | 5 | 14.18 | 0.000621 |
| Leishmanolysin (peptidase family M8) [PF01457.19] | 6 | 5 | 14.18 | 0.000621 |
| TM(4x) | 415 | 45 | 1.84 | 0.000664 |
| Potassium channel inhibitor ShK domain-like [PF01549.27] | 111 | 19 | 2.91 | 0.00367 |
| Immunoglobulin C1-set domain [PF07654.18] | 12 | 6 | 8.51 | 0.00397 |
| CD80-like C2-set immunoglobulin domain [PF08205.15] | 29 | 9 | 5.28 | 0.00397 |
| Myosin N-terminal SH3-like domain [PF02736.22] | 8 | 5 | 10.63 | 0.00461 |
| Sodium:neurotransmitter symporter family (SNF) [PF00209.21] | 24 | 8 | 5.67 | 0.00587 |
| Peptidase family M13/neprilysin metallopeptidase [PF01431.24] | 39 | 10 | 4.36 | 0.00819 |

This table shows 10 Pfam protein motifs and four overrepresented signal or transmembrane domain configurations predicted by Phobius that are overrepresented in the positively immunoregulated intestinal gene set (1,951 out of 33,190 genes) with q-values of $\leq 0.01$ . Phobius domains are abbreviated with 'SigP'

Genes with types of immunoregulation other than positive intestinal had fewer notable protein families (Supplementary Table S15). The 137 negatively immunoregulated intestinal genes disproportionately encoded cysteine proteases and saposins, with saposins being uniquely enriched in this gene set (36-fold over chance; q = 9.94•10^−3^). At least four *C. elegans* saposins have antimicrobial activity *in vitro* (Banyai and Patthy 1998; Roeder *et al*. 2010; Hoeckendorf and Leippe 2012; Hoeckendorf *et al*. 2012); conversely, one saposin in *A. ceylanicum* has no antibacterial activity but does lyse blood cells *in vitro* (He *et al*. 2021). Negatively immunoregulated hookworm saposins might have either or both functions. Of the 26 positively immunoregulated non-intestinal genes, 15 encoded ASP proteins with CAP domains (46-fold over chance; q = 7.61•10^−19^); 10 had non-intestine-biased expression (10.7-fold over chance; p = 1.12•10^−8^). Both gene sets disproportionately encoded secreted proteins; neither set had prominently sex-biased expression.

### Immunoregulated genes have significantly male-biased expression

Multicellular parasites such as hookworms are generally studied to understand mechanisms of infection common to both sexes, and indeed we ourselves collected both ES proteins and RNA from mixed-sex populations. However, males of the African tick *Rhipicephalus appendiculatus* excrete immunoglobulin-binding proteins that the males themselves do not seem to need, and that are instead required for coinfecting female ticks to feed on host blood efficiently (Wang *et al*. 1998). This instance of sexual cooperation in a parasite made us wonder whether there existed male-biased or female-biased genes among our immunoregulated gene set. Using published RNA-seq data from *A. ceylanicum* adult males and females (Bernot *et al*. 2020), we observed that 3,135 and 1,547 of our 33,190 predicted *A. ceylanicum* genes had male- or female-biased expression, respectively (Table 6). Going on to check for overlaps of these gene sets with our immunoregulated gene set, we found that 50.1% of our positively immunoregulated intestinal genes (977/1,951; 5.3-fold over background; p = 0) were also male-biased, while only 11.6% of them (227/1,951; 2.5-fold over background; p = 3.34•10^−38^) were female-biased (Table 9; Supplementary Table S15).

**Table 9.** Gene categories overrepresented in *A. ceylanicum* immunoregulated genes.

| **Immunoreg. category** | **Other category** | **Immunoreg. genes** | **Other genes** | **Overlap** | **Enrichment** | **p-value** |
| --- | --- | --- | --- | --- | --- | --- |
| Pos. intestinal | Male-biased | 1,951 | 3,135 | 977 | 5.30 | 0 |
| Pos. intestinal | Female-biased | 1,951 | 1,547 | 227 | 2.50 | 3.34e-38 |
| Pos. intestinal | Intestine-biased | 1,951 | 1,670 | 1 | 0.0102 | 1.54e-43 |
| Pos. intestinal | Non-intestine-biased | 1,951 | 1,196 | 258 | 3.67 | 5.51e-78 |
| Pos. intestinal | All ES | 1,951 | 565 | 153 | 4.61 | 1.19e-59 |
| Neg. intestinal | Male-biased | 137 | 3,135 | 20 | 1.55 | 0.0548 |
| Neg. intestinal | Female-biased | 137 | 1,547 | 2 | 0.31 | 0.0988 |
| Neg. intestinal | Intestine-biased | 137 | 1,670 | 50 | 7.25 | 6.56e-30 |
| Neg. intestinal | Non-intestine-biased | 137 | 1,196 | 4 | 0.810 | 1 |
| Neg. intestinal | All ES | 137 | 565 | 8 | 3.43 | 0.00247 |
| Pos.non-intestinal | Male-biased | 26 | 3,135 | 6 | 2.44 | 0.0312 |
| Pos.non-intestinal | Female-biased | 26 | 1,547 | 0 | 0 | 0.632 |
| Pos.non-intestinal | Intestine-biased | 26 | 1,670 | 0 | 0 | 0.640 |
| Pos.non-intestinal | Non-intestine-biased | 26 | 1,196 | 10 | 10.67 | 1.12e-08 |
| Pos.non-intestinal | All ES | 26 | 565 | 13 | 29.37 | 7.47e-17 |
| Pos. int. or non-int. | All ES | 1,975 | 565 | 164 | 4.88 | 5.53e-68 |

Such predominantly male-biased expression, along with the enrichment of three male-associated gene families (MSP, MFP2, and DUF236) in this gene set, raises the question of whether the changes in hookworm intestinal gene expression we observed with different host immunological backgrounds were actually due to our having harvested a higher proportion of male hookworms from normal than immunosuppressed hosts, leading to a male skew in putatively immunoregulated genes. We see two reasons why such bias is not sufficient to explain our observations. First, we only observed significant changes of gene activity in dissected intestines from 19-day adults, while observing no such changes in non-intestinal tissues from those same adults (Table 6). If apparent immunoregulation of genes had actually been due to overlooked male bias in the hookworms we collected from normal hosts, one would expect to see at least as much (if not more) changed gene expression in non-intestinal tissues (which contained most, if not all, of the gonadal tissue from adults) as we saw from dissected intestines. The absence of significant immunoregulation in non-intestinal tissues (or in our 12-day whole animals) is inconsistent with such an artifact. Second, positively immunoregulated intestinal genes are not a simple subset of male-biased genes, as would be expected if male selection accounted for immunological changes of gene expression: 747 of these 1,951 immunoregulated genes had no sex bias, and 227 of them had female-biased expression. We conclude that the transcriptional immunoregulation we describe here, though dominated by male-biased genes, is nevertheless real. This suggests that male and female *A. ceylanicum* have different responses to the host immune system, and that such differences might account for the observed male-bias from our mixed-sex *A. ceylanicum* 19-day intestinal RNA-seq data.

### The host immune system affects ES gene expression

Protein products of ES genes are thought to affect the host immune system, but whether the host immune system affects ES genes has been unclear. Comparing the ES gene set with immunomodulated gene sets, we observed significant overlaps (Table 9). Out of 1,951 positively immunoregulated intestinal genes in 19-day hookworms, 153 also encoded ES proteins (27.1% of ES genes; 4.6-fold over chance; p = 1.19•10^−59^). Of 26 positively immunoregulated non-intestinal genes, 13 were also ES genes (29-fold above chance; p = 7.47•10^−17^); such expression might reflect synthesis and secretion of ES proteins by cephalic/pharyngeal glands (Huang *et al*. 2020).

The 153 positively immunoregulated intestinal ES genes disproportionately encoded astacin proteases, TIMP and TIL protease inhibitors, ShK-like proteins, ASPs, ASPRs, and SCVPs (Table 10). Of these genes, 69 had ES gene homologs in *A. caninum* (20-fold enrichment; p = 6.00•10^−72^), 24 in *N. americanus* (11-fold enrichment; p = 2.35•10^−18^), and 24 in *H. contortus* (4.4-fold enrichment; p = 1.23•10^−9^; Supplementary Table S16). As noted above, all of the above protein families may affect immunomodulation in hookworm hosts; in addition, astacin has been shown to enable tissue invasion *in vitro* by *A. caninum* (Williamson *et al*. 2006). Other immunoregulated ES genes also encoded proteins with possible functions in immunoregulation, antithrombosis, or digesting host tissue (Supplementary Table S6): the ASPs *Acey*-NIF-B (Moyle *et al*. 1994; Lo *et al*. 1999) and *Acey*-HPI (Del Valle *et al*. 2003), two apyrases (Guiguet *et al*. 2016; Pala *et al*. 2023), deoxyribonuclease II (Bouchery *et al*. 2020), hyaluronidase (Hotez *et al*. 1992; Yang *et al*. 2020), and superoxide dismutase (Knox and Jones 1992; Brophy *et al*. 1995).

**Table 10.**
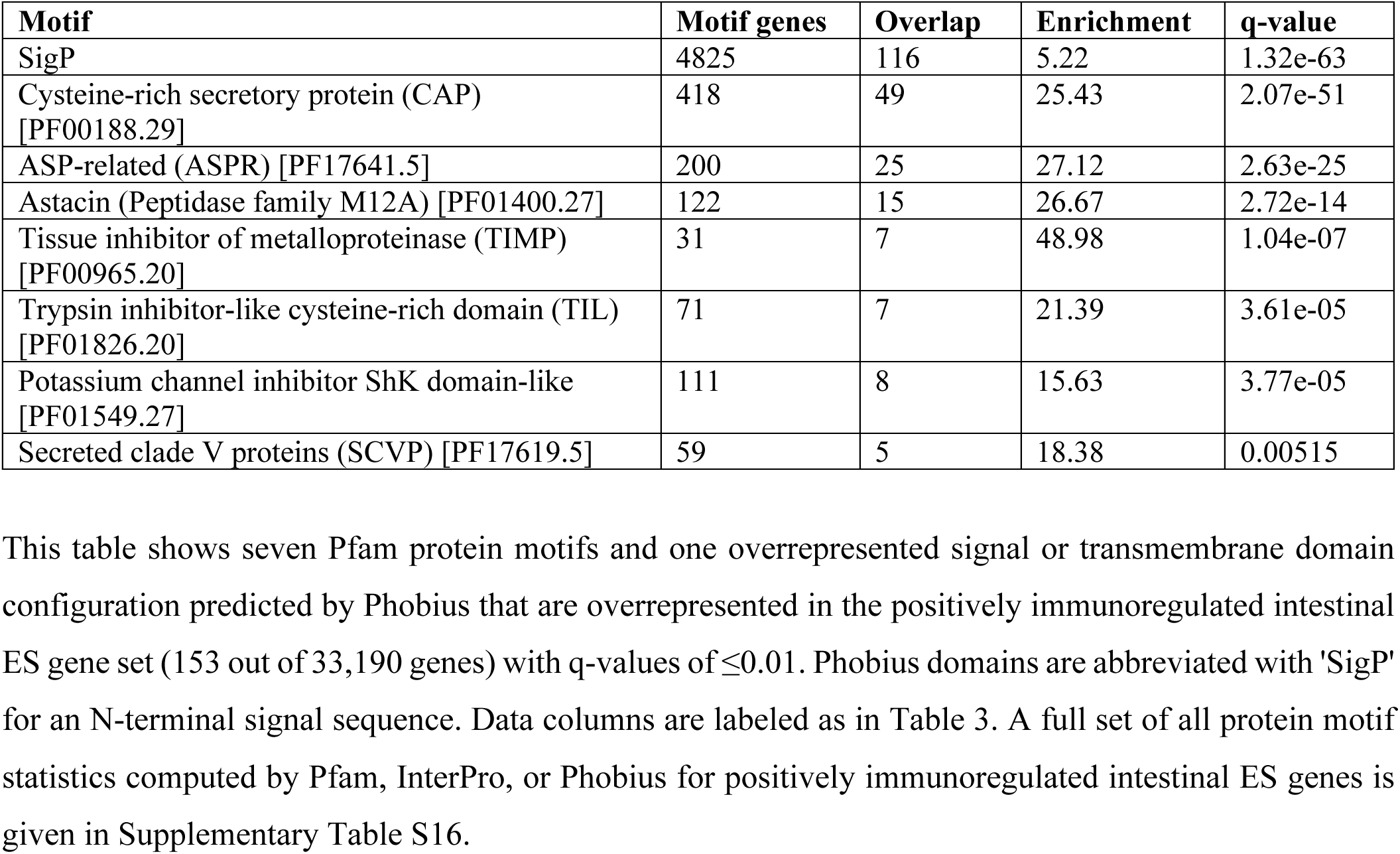
Protein motifs overrepresented in positively immunoregulated intestinal *A. ceylanicum* ES genes.

| **Motif** | **Motif genes** | **Overlap** | **Enrichment** | **q-value** |
| --- | --- | --- | --- | --- |
| SigP | 4825 | 116 | 5.22 | 1.32e-63 |
| Cysteine-rich secretory protein (CAP) [PF00188.29] | 418 | 49 | 25.43 | 2.07e-51 |
| ASP-related (ASPR) [PF17641.5] | 200 | 25 | 27.12 | 2.63e-25 |
| Astacin (Peptidase family M12A) [PF01400.27] | 122 | 15 | 26.67 | 2.72e-14 |
| Tissue inhibitor of metalloproteinase (TIMP) [PF00965.20] | 31 | 7 | 48.98 | 1.04e-07 |
| Trypsin inhibitor-like cysteine-rich domain (TIL) [PF01826.20] | 71 | 7 | 21.39 | 3.61e-05 |
| Potassium channel inhibitor ShK domain-like [PF01549.27] | 111 | 8 | 15.63 | 3.77e-05 |
| Secreted clade V proteins (SCVP) [PF17619.5] | 59 | 5 | 18.38 | 0.00515 |

## Discussion

The ability of hookworms to suppress or evade their hosts’ immune systems and feed on their hosts’ blood and other tissues, for years, has motivated identifying hookworm genes whose products interact with the host, either by encoding ES proteins or by directly contacting host blood and tissues in the hookworm intestine (Wei *et al*. 2016; Abuzeid *et al*. 2020). Here we have identified ES proteins, intestine-biased genes, and immunoregulated genes in the hookworm *A. ceylanicum*, the last of which is unique to this study. Indeed, we hypothesized that hookworms regulate genes in response to the host immune system in order to suppress it, which motivated our immunosuppression/transcriptomic study here. Strongylid parasitic nematodes interact with the immune system despite generally not being blood-feeding (Bansemir and Sukhdeo 1994; McNeilly *et al*. 2017; Price *et al*. 2019), so blood feeding is not necessary for such interactions. However, in the case of hookworms, our findings imply that interaction with the host immune system primarily occurs through contact of the hookworm intestinal lumen with host blood.

Zoonotic *A. ceylanicum* productively infects both humans and other mammals, making it important to humans while also enabling laboratory studies of a true hookworm that is clinically relevant (Traub 2013; Colella *et al*. 2021). We have analyzed both our new gene sets and previously identified gene sets (Wei *et al*. 2016; Bernot *et al*. 2020; Uzoechi *et al*. 2023), to identify traits encoded either by ES genes, intestinal genes, immuoregulated genes, or all three. To enable this work, we repredicted protein-coding genes in *A. ceylanicum* to the highest completeness so far achieved. We have greatly expanded the number of intestine-biased genes in *A. ceylanicum* and found them to be extensively homologous to intestine-biased genes in *H. contortus* (Laing *et al*. 2013). We identified a mixture of functionally suggestive and poorly understood protein families in *A. ceylanicum* ES proteins and intestine-biased genes, observed genes that are immunoregulated in adult intestines (where they are exposed to host blood and circulating immune factors) but not other tissues, detected an unexpected male bias in positively immunoregulated intestinal genes, and defined a positively immunoregulated subset of ES genes that have a mosaic of conservation and species-specificity.

On examination, it was possible to infer biologically coherent functions from what might seem to be a jumble of hookworm ES proteins. Immunosuppressors have been long suspected to be part of the ES protein repertoire, and we observed several protein types that could have this role. We also observed a pattern of diverse multigene families encoding ES proteins (ASPs, ASPRs, TTLs, and SCVPs); in other parasitic nematodes, such a pattern has been repeatedly seen both in ES proteins and in genetic diversity between strains or sibling species (Abuzeid *et al*. 2020; Cole *et al*. 2023; Stevens *et al*. 2023). It is not clear why this pattern exists; perhaps hookworms and other parasitic nematodes use diverse ES multigene families to survive unpredictably varying host immune systems by confusing their hosts with complex and varying mixtures of protein antigens, which leads parasitic nematodes to accumulate multigene families through balancing selection (Thomas and Robertson 2008; Tellier *et al*. 2014; Ebert and Fields 2020). Another possible function that consistently emerged when examining ES proteins was inhibition of blood clotting. Despite being less often discussed and less obviously virulent, antithrombotics are at least as important for parasitism as immunosuppressants. Hookworm anticoagulants were first observed in 1904 (Loeb and Smith 1904), and are likely to be necessary for sustained blood-feeding (Kvist *et al*. 2020; Oliveira and Genta 2021; Pala *et al*. 2023).

The strong overlap that we observed between immunoregulation and male-biased gene expression raises the question of whether hookworms have sex-specialized repertoires of genes that they express during infections (Costa *et al*. 2009). There has been one striking instance (in the African tick *R. appendiculatus*) where males and females interact differently with their hosts and the males act to promote female success during parasitism (Wang *et al*. 1998). Our current data do not indicate whether such cooperation happens during hookworm infections, let alone whether the male-biased immunoregulated genes we describe are relevant to such hypothetical cooperation. However, such a bias might ensure that both sexes are present in a successful infection, which is required for reproduction. More extensive analysis to test for sexual cooperation during infections or sex-biased gene immunoregulation should be pursued for hookworms and other parasitic nematodes.

Immunoregulated genes of *A. ceylanicum* have homologs in *A. caninum*, *N. americanus*, and *H. contortus*. The homologs of immunoregulated ES genes also encode ES proteins, suggesting that other parasitic nematodes may also have immunoregulated genes, and some findings indicate that they actually do. For the strongylid *Teladorsagia circumcincta*, being embedded in host mucosa upregulates ES genes and putative immunomodulatory genes (McNeilly *et al*. 2017; Price *et al*. 2019). For the strongylid *H. bakeri*, infecting mouse hosts with colitis induces different ES proteins than normal hosts (Maruszewska-Cheruiyot *et al*. 2023). Finally, for the strongylid *N. brasiliensis*, recent transcriptomic and proteomic analyses shows genes that are positively or negatively regulated during infections of normal versus immunocompromised *stat6* mutant mouse hosts, a subset of which encode ES proteins (Ferguson *et al*. 2023; Ferguson *et al*. 2024). Thus, the immunoregulation we observe here for *A. ceylanicum* may be a general phenomenon found in many other parasitic nematodes.

## Supplementary Table, Figure, and File Legends

**Supplementary Table S1.** *A. ceylanicum* RNA-seq libraries newly generated in this study. “RNA-seq library ID” provides a short abbreviation for each library; these abbreviations have been used for expression levels (TPMs), mapped read counts (reads), and differential gene expression analyses in later supplementary data tables. For each library, “SRA accession”, “BioProject accession” and “BioSample accession” provide accession numbers; “Reads”, “Total nt”, and “Read length in nt” give the number of reads, total sequence in nt, and read length; and “RNA-seq description” summarizes biological contents.

**Supplementary Table S2.** Software used in this study. “Software” provides the name of each computer program or suite of computer programs; “Purpose” describes why this software was used here; “Main web site (URL) or code location” gives the primary Web site for this software; “Bioconda source (if used)” gives the bioconda web site for programs that were installed as bioconda environments; “Online documentation” gives the web site for detailed online manuals, if a given program has one. Publications and arguments for each program are cited and described in Methods.

**Supplementary Table S3.** Published genomic data used in this study. Genome sequences, coding sequences, proteomes, and gene annotations of relevant nematodes were downloaded from WormBase (Davis *et al*. 2022) or ParaSite (Howe *et al*. 2017; Lee *et al*. 2017) and used for transcriptomic or proteomic analyses; they include Coghlan gene predictions and their corresponding *A. ceylanicum* genome assembly, along with UniProt (UniProt 2023) and RefSeq (O’Leary *et al*. 2016) proteome databases from highly GO-annotated model organisms. Four separate spreadsheets are given for data files of genomes, proteomes, CDS DNA sets, and gene annotations in GFF format. For each data file, “Species” gives the biological species described by the file, “Comments” describes the particular use(s) to which the data file was put in this study, and “URL” gives the Web source of the file.

**Supplementary Table S4.** Published RNA-seq data of *A. ceylanicum* (Schwarz *et al*. 2015; Wei *et al*. 2016; Bernot *et al*. 2020) and *H. contortus* (Laing *et al*. 2013) used in this study. All RNA-seq files were downloaded from the European Nucleotide Archive (ENA). “Species” gives the file’s origin species. “Abbreviation” provides a short abbreviation for each library; these abbreviations have been used for expression levels (TPMs), mapped read counts (reads), and differential gene expression analyses in later supplementary data tables. “Biological condition (sex, developmental stage, tissue) and replicate number” describes biological content and replication. “Notes” describes any ambiguities that had to be resolved during analysis of the data. “Database” is uniformly ‘ENA’, but is included for completeness in the table. “Accession” and “URL” give SRA accession numbers and ENA Web sources for each file.

**Supplementary Table S5.** LC-MS/MS analyses of two *A. ceylanicum* ES protein sets, E20201108-05 and E20201108-07. Data columns were generated by Proteome Discoverer 2.4; other details of the analysis are given in Methods.

**Supplementary Table S6.** Annotations, RNA-seq expression, and differential gene expression for the *A. ceylanicum* v2.1 protein-coding gene set. Its data columns are as follows.

<u>Gene identifications and equivalencies</u>

**Gene:** A given predicted protein-coding gene in the *A. ceylanicum* genome assembly. All further data columns are pertinent to that particular gene.

**Gene_name:** A human-readable gene name, where it exists (e.g., “*Acey*-CP-1” instead of simply “Acey_s0154.v2.g3234”). These names are used in the main text to discuss genes of particular interest.

**Mapped_v1.0_gene:** Any gene or genes from our previous v1.0 predictions which overlap the v2.1 gene here by least 1 nt of coding exon sequence in the *A. ceylanicum* genome. Although this criterion errs on the side of sensitivity and we have made no effort to filter out short overlaps, we expect that in practice this will identify extensive exon overlaps between an earlier v1.0 gene and its reprediction in v2.1.

**Mapped_Coghlan_gene:** Any gene or genes from the previous *A. ceylanicum* gene predictions by Coghlan *et al*. (Coghlan *et al*. 2019) which, when lifted over onto our *A. ceylanicum* genome assembly, overlap the v2.1 gene here by least 1 nt of coding exon sequence.

**Mapped_Uzoechi_gene:** Any gene or genes from the recent *A. ceylanicum* gene repredictions by Uzoechi *et al*. (Uzoechi *et al*. 2023) which overlap the v2.1 gene here by least 1 nt of coding exon sequence.

*General traits of protein products*

**Max_prot_size:** The size of the largest predicted protein product.

**Prot_size:** This shows the full range of sizes for all protein products from a gene’s predicted isoforms.

**Phobius:** This denotes predictions of signal and transmembrane sequences made with Phobius 1.01 (KÄll *et al*. 2004). ‘SigP’ indicates a predicted signal sequence, and ‘TM’ indicates one or more transmembrane-spanning helices, with N helices indicated with ‘(Nx)’. Varying predictions from different isoforms are listed.

**NCoils:** This shows coiled-coil domains, predicted by Ncoils 2002.08.22 (Lupas 1996). Both the proportion of such sequence (ranging from 0.01 to 1.00) and the exact ratio of coiled residues to total residues are given. Proteins with no predicted coiled residues are blank.

**Psegs:** This shows what fraction of a protein is low-complexity sequence, as detected by PSEG 1999.06.10 (Wootton 1994). As with Ncoils, relative and absolute fractions of low-complexity residues are shown.

**Pfam:** Predicted protein domains from Pfam 35.0 (Mistry *et al*. 2021), with family-specific significance thresholds.

**InterPro:** Predicted protein domains from InterProScan 5.57-90.0 (Paysan-Lafosse *et al*. 2023).

**AMP:** An antimicrobial peptide (AMP) gene annotation for v1.0 of our *A. ceylanicum* gene predictions, predicted by Irvine *et al*. (Irvine *et al*. 2023), and mapped onto our v2.1 gene predictions here.

**GO_Biological:** Annotations from the biological subset of Gene Ontology (GO) terms (Ashburner *et al*. 2000; Carbon *et al*. 2021), generated with EnTAP 0.10.7-beta (Hart *et al*. 2020).

**GO_Molecular:** Annotations from the molecular subset of Gene Ontology (GO) terms (Ashburner *et al*. 2000; Carbon *et al*. 2021), generated with EnTAP 0.10.7-beta (Hart *et al*. 2020).

**GO_Cellular:** Annotations from the cellular subset of Gene Ontology (GO) terms (Ashburner *et al*. 2000; Carbon *et al*. 2021), generated with EnTAP 0.10.7-beta (Hart *et al*. 2020).

**EggNOG_description:** EggNOG descriptions (Hernandez-Plaza *et al*. 2023), generated with EnTAP 0.10.7-beta (Hart *et al*. 2020).

**EggNOG_KEGG:** KEGG codes (Kanehisa *et al*. 2023), generated with EnTAP 0.10.7-beta (Hart *et al*. 2020).

<u>Orthologies of protein products</u>

**OFind_Summary** and **OFind_Full**

The results for our OrthoFinder analysis (Emms and Kelly 2019) of orthologies between *A. ceylanicum* and the related nematodes *A. caninum*, *C. elegans*, *H. contortus*, *H. bakeri*, *N. americanus*, *N. brasiliensis*, *Pristionchus pacificus*, and *T. circumcincta*. For one of these species (*N. americanus*) two different proteomes were included: the original predicted proteome by Tang *et al*. (Tang *et al*. 2014) labeled ‘necator_orig’, and the repredicted proteome by Logan *et al*. (Logan *et al*. 2020) labeled ‘necator_rev’. Two different views of these results are given: the summary lists taxa and gene counts, while the full results give individual gene names.

<u>ES- and gene-expression-related traits</u>

**Intest_haemonchus:** Homologous *H. contortus* genes (taken from **OFind_Full**) that have intestine-biased gene expression, as computed by our analysis of previously published *H. contortus* RNA-seq data (Supplementary Table S4).

**Non-intest_haemonchus:** Homologous *H. contortus* genes (taken from **OFind_Full**) that have non-intestine-biased gene expression, as computed by our analysis of previously published *H. contortus* RNA-seq data (Supplementary Table S4).

**ES_component:** A gene whose protein product was detected in at least one of our two ES mass spectrometry experiments, E20201108-05 and E20201108-07. Note that this was used as a binary (Boolean) classification for statistical analyses of gene set overlaps; see below for other such binary classifications.

**ES_observations:** Observations of a gene’s protein product being present in either E20201108-05, or E20201108-07, or both.

**Coghlan-spec_ES:** In our mass spectrometry analysis, our observation of a peptide mapping specifically to a Coghlan gene-encoded protein that was distinct from our v2.1 proteins by at least one amino acid residue, but whose Coghlan gene’s coding exons could then be lifted over to the coding exons of the v2.1 gene here (**Mapped_Coghlan_gene**). For each such mapping, the ES observations behind it are noted (either E20201108-05, or E20201108-07, or both).

**Uzoechi_ES:** An ES-encoding gene detected by Uzoechi *et al*. (Uzoechi *et al*. 2023) whose coding exons mapped onto the coding exons of this v2.1 gene (in **Mapped_Uzoechi_gene**). Note that Uzoechi *et al*. separately observed both male and female ES proteins, and those specific observations are given here.

**ES_a_caninum:** Homology to an *A. caninum* gene encoding an *A. caninum* ES protein observed by Morante *et al*. (Morante *et al*. 2017), extracted from **OFind_Full**.

**ES_necator_rev:** Homology to an *N. americanus* gene encoding an *N. americanus* ES protein observed by Logan *et al*. (Logan *et al*. 2020), extracted from **OFind_Full**.

**ES_haemonchus:** Homology to an *H. contortus* gene encoding an *H. contortus* ES protein observed by Wang *et al*. (Wang *et al*. 2019), extracted from **OFind_Full**.

<u>Binary (Boolean) classifications of genes</u>

**Male-biased:** Annotation here indicates a gene with male-biased gene expression, as defined by ≥2-fold higher expression in males versus females, with a false discovery rate (FDR) of ≤ 0.01.

**Female-biased:** Annotation here indicates a gene with female-biased gene expression, as defined by ≥2-fold higher expression in females versus males, with a false discovery rate (FDR) of ≤ 0.01.

**Intestine-biased:** Annotation here indicates a gene with intestine-biased gene expression, as defined by ≥2-fold higher expression in 19-day intestinal tissue from normal hosts versus 19-day non-intestinal tissue from normal hosts, with a false discovery rate (FDR) of ≤ 0.01.

**Non-intestine-biased:** Annotation here indicates a gene with non-intestine-biased gene expression, as defined by ≥2-fold higher expression in 19-day non-intestinal tissue from normal hosts versus 19-day intestinal tissue from normal hosts, with a false discovery rate (FDR) of ≤ 0.01.

**Any.19d.immunoreg:** Annotation here indicates a gene with either intestinal or non-intestinal and either positively or negatively immunoregulated gene expression, as defined by ≥2-fold higher or lower expression in 19-day intestinal or non-intestinal tissue from normal hosts versus 19-day corresponding tissue from dexamethasone-immunosuppressed hosts, with a false discovery rate (FDR) of ≤ 0.01.

**Any.19d.pos.immunoreg:** Annotation here indicates a gene with either intestinal or non-intestinal positively immunoregulated gene expression, as defined by ≥2-fold higher in 19-day intestinal or non-intestinal tissue from normal hosts versus 19-day corresponding tissue from dexamethasone-immunosuppressed hosts, with a false discovery rate (FDR) of ≤ 0.01.

**Pos.intest.immunoreg:** Annotation here indicates a gene with intestinal positively immunoregulated gene expression, as defined by ≥2-fold higher expression in 19-day intestinal tissue from normal hosts versus 19-day intestinal tissue from dexamethasone-immunosuppressed hosts, with a false discovery rate (FDR) of ≤ 0.01.

**Neg.intest.immunoreg:** Annotation here indicates a gene with intestinal negatively immunoregulated gene expression, as defined by ≥2-fold higher expression in 19-day intestinal tissue from dexamethasone-immunosuppressed hosts versus 19-day intestinal tissue from normal hosts, with a false discovery rate (FDR) of ≤ 0.01.

**Pos.non-intest.immunoreg:** Annotation here indicates a gene with non-intestinal positively immunoregulated gene expression, as defined by ≥2-fold higher expression in 19-day non-intestinal tissue from normal hosts versus 19-day non-intestinal tissue from dexamethasone-immunosuppressed hosts, with a false discovery rate (FDR) of ≤ 0.01.

**Neg.non-intest.immunoreg:** Annotation here indicates a gene with non-intestinal negatively immunoregulated gene expression, as defined by ≥2-fold higher expression in 19-day non-intestinal tissue from dexamethasone-immunosuppressed hosts versus 19-day non-intestinal tissue from normal hosts, with a false discovery rate (FDR) of ≤ 0.01.

**Neg.12d.immunoreg:** Annotation here indicates a gene with negatively immunoregulated gene expression in 12-day young adults, as defined by ≥2-fold higher expression in 12-day young adults from dexamethasone-immunosuppressed hosts versus 12-day young adults from normal hosts, with a false discovery rate (FDR) of ≤ 0.01. (Note that no genes were observed with positively immunoregulated gene expression in 12-day young adults, so no ‘**Pos.12d.immunoreg**’ data column was needed.)

**Uzoechi_any_ES:** Annotation here indicates a gene which was detected as encoding some sort of ES protein by Uzoechi *et al*. (Uzoechi *et al*. 2023). Practically, this means a gene with any annotation in **Uzoechi_ES** as encoding either a male ES (’Uzoechi_male_ES’), or a female ES (’Uzoechi_female_ES’), or both.

**Uzoechi_male.only_ES:** Annotation here indicates a gene which was detected as encoding a male-specific ES protein by Uzoechi *et al*. (Uzoechi *et al*. 2023), annotated as solely ‘Uzoechi_male_ES’ in **Uzoechi_ES**.

**Uzoechi_female.only_ES:** Annotation here indicates a gene which was detected as encoding a female-specific ES protein by Uzoechi *et al*. (Uzoechi *et al*. 2023), annotated as solely ‘Uzoechi_female_ES’ in **Uzoechi_ES**.

**Uzoechi_both_ES:** Annotation here indicates a gene which was detected as encoding an ES protein in both males and females by Uzoechi *et al*. (Uzoechi *et al*. 2023), annotated as both ‘Uzoechi_male_ES’ and ‘Uzoechi_female_ES’ in **Uzoechi_ES**.

<u>Gene expression</u>

**[X].TPM:** For each individual RNA-seq data set (with ‘X’ denoting the data set’s abbreviation), this gives gene expression levels in TPM, computed by Salmon 1.9.0 (Patro *et al*. 2017). Keys to all abbreviations are given in Supplementary Tables Sx and Sy. Biological replicates of RNA-seq samples are denoted by suffixes such as ‘_1’, ‘_2’, or ‘_3’.

**[X].reads:** For each individual RNA-seq data set (with ‘X’ denoting the data set’s abbreviation), this gives numbers of mapped RNA-seq reads per gene, computed for individual RNA-seq data sets by Salmon 1.9.0 (Patro *et al*. 2017), with fractional values rounded down to integers.

<u>Differential gene expression between biological conditions</u>

**[X].vs.[Y].logFC:** The fold-changes of gene expression between biological condition X and biological condition Y, expressed as log2 values, and with positive values representing greater expression in condition X. The values listed here are only those were computed to be significant using edgeR 3.36.0 (Robinson *et al*. 2010), with multiple biological RNA-seq replicates for most conditions being compared, with all biological replicates being analyzed in a single edgeR run, and with significant results annotated for individual genes. Biological conditions of RNA-seq samples are abbreviated as shown in **[X].TPM** or **[X].reads** but without the replicate suffixes.

**[X].vs.[Y].FDR:** The false discovery rate (FDR) for gene expression changes between biological condition X and biological condition Y, annotated for individual genes. The FDR for a given set of positive results is defined as that significance threshold which, if accepted, will lead to the entire set of positives having a collective false-positive rate no greater than the FDR; it therefore provides a way to correct for testing multiple hypotheses without rejecting excessive numbers of true positives. As with **[X].vs.[Y].logFC**, only changes that were computed to be significant by edgeR are listed.

**Supplementary Table S7.** Statistical analyses of the overlaps between *A. ceylanicum* genes encoding protein motifs (or other traits) and *A. ceylanicum* genes encoding ES proteins. Each category of genes (e.g., genes encoding a particular protein motif or having a particular trait) was compared for its degree of overlap to a set of ES genes, and statistically analyzed for the non-randomness of this overlap by a two-tailed Fisher test. In situtations where many categories were compared at once (for instance, an entire set of protein motifs from Pfam, InterPro, or Phobius), p-values initially generated by Fisher testing were used to compute q-value significance scores which corrected for multiple hypothesis testing. Statistics are provided both for the set of 565 ES genes described in this paper (labeled “UMass_ES”) and for the set of 860 ES genes described by Uzoechi *et al*. (Uzoechi *et al*. 2023) (labeled “WashU_ES”). For each ES set, spreadsheets are given for analyses of overlaps with the following traits; signal or transmembrane domain organizations predicted by Phobius; protein motifs in Pfam; protein motifs in InterPro; and various binary comparisons with other gene traits (“Various”). Each analysis provides the following data: “Motif” denotes the specific protein motif (or other gene category) for which genes encoding it are being tested for nonrandomly high (or low) overlap with the ES gene set; “All_genes” gives the total number of v2.1 protein-coding genes within which overlaps were tested; “Motif_genes” gives the number of genes annotated with the protein motif (or other trait) being tested for overlap; “Class_genes” gives the number of ES genes being tested for overlap; “Motif.Class_overlap” gives the observed number of genes falling into both categories; “Exp_rand_overlap” gives the number of genes that would be expected to overlap purely randomly; “Enrichment” gives the ratio of observed versus random overlaps (note that this ratio can be lower than 1, and in fact can be as low as 0); “p-value” gives an initial stastistical significance for the observed overlap, computed by a two-tailed Fisher test; “q-value” gives, for cases of many comparisons at once (e.g., testing for all Pfam motifs simultaneously), a significance score that conservatively corrects for multiple hypothesis testing (Bailey *et al*. 2009; Noble 2009). Note that q-values were not computed for simple binary comparisons of gene traits listed in “Various”, which are annotated for *A. ceylanicum* genes in **Supplementary Table S6**, and defined in its table legend: Intestine-biased; Non-intestine-biased; Pos.intest.immunoreg; Neg.intest.immunoreg; Pos.non-intest.immunoreg; Neg.non-intest.immunoreg; Male-biased; Female-biased; Uzoechi_any_ES; Uzoechi_male.only_ES; Uzoechi_female.only_ES; and Uzoechi_both_ES. Also, again note that highly significant overlaps can be either higher or lower than the randomly expected genome-wide background rate.

**Supplementary Table S8.** Annotations, RNA-seq expression, and differential gene expresssion for the *H. contortus* protein-coding gene set. We annotated predicted protein products with N-terminal signal sequences, conserved protein domains, orthologies to genes in related nematode species, and Gene Ontology (GO) terms describing biological and molecular functions. Protein-coding genes were predicted by Doyle *et al*. (Doyle *et al*. 2020). Since the annotation methods we used for this *H. contortus* proteome were identical to those we used for our *A. ceylanicum* v2.1 proteome, almost all the data columns used here are equivalent to those in **Supplementary Table S6**. One data column unique to this table is “Hco_ES”, which denotes *H. contortus* genes previously demonstrated by Wang *et al*. (Wang *et al*. 2019) to encode ES proteins.

**Supplementary Table S9.** Annotations for protein-coding gene sets of *A. caninum* and *N. americanus*. We annotated predicted protein products with N-terminal signal sequences, conserved protein domains, orthologies to genes in related nematode species, and Gene Ontology (GO) terms describing biological and molecular functions. Protein-coding genes were predicted for *A. caninum* by Coghlan *et al*. (Coghlan *et al*. 2019) and for *N. americanus* by Logan *et al*. (Logan *et al*. 2020). Since the annotation methods we used for these proteomes were identical to those we used for our *A. ceylanicum* v2.1 proteome, almost all the data columns used here are equivalent to those in **Supplementary Table S6**. Two data columns unique to these tables are **Acan_ES** and **Necator_ES**, which respectively denote *A. caninum* or *N. americanus* genes previously demonstrated by Morante *et al*. (Morante *et al*. 2017) or Logan *et al*. (Logan *et al*. 2020) to encode ES proteins.

**Supplementary Table S10.** Statistical analyses of the overlaps between *A. caninum*, *N. americanus*, or *H. contortus* genes encoding protein motifs (or other traits) and *A. caninum*, *N. americanus*, or *H. contortus* genes encoding ES proteins. ES gene annotations are taken from Supplementary Tables S8 and S9. Statistical significances of overlaps between motif/trait genes and ES genes were computed and described as in **Supplementary Table S7**.

**Supplementary Table S11.** Statistical analyses of overlaps between pairs of *A. ceylanicum* gene sets, with *A. ceylanicum* genes encoding ES proteins versus *A. ceylanicum* genes having various homologies to either ES genes or any genes in *A. caninum*, *H. contortus*, or *N. americanus*. “ES_a_caninum”, “ES_haemonchus”, and “ES_necator_rev” denote sets of *A. ceylanicum* genes with homology to ES genes in *A. caninum*, *H. contortus*, or *N. americanus*; “ES_Acan.or.Hco.or.L2020” denotes a set of *A. ceylanicum* genes with homology to ES genes in any of these three species. “Acan_homology”, “Haemonchus_homology”, and “Necator_homology” denote sets of *A. ceylanicum* genes with homology to any genes (ES protein-encoding, or not) in *A. caninum*, *H. contortus*, or *N. americanus*; “Acan.Hco.Nec_homology” denotes a set of *A. ceylanicum* genes with homology to any genes in any of these three species. The ES gene sets are either from this study (“all_UMass_ES”) or from Uzoechi *et al*. (Uzoechi *et al*. 2023) (“all_WashU_ES”). Subsets of these ES gene sets that lack homologies to known ES genes in related parasites are denoted with “non_AcanES_homol_[ES]” (for *A. ceylanicum* ES genes lacking homologies to *A. caninum* ES genes), “non_HcoES_homol_[ES]” (for *A. ceylanicum* ES genes lacking homologies to *H. contortus* ES genes), “non_NecES_homol_[ES]” (for *A. ceylanicum* ES genes lacking homologies to *N. americanus* ES genes), or “non_anyES_homol_[ES]” (for *A. ceylanicum* ES genes lacking homologies to ES genes in any of the three other species). Statistical significances of overlaps between homologous genes and ES genes were computed and described as in **Supplementary Table S7**.

**Supplementary Table S12**. The frequencies with which RNA-seq reads (either from our new RNA-seq libraries listed in **Supplementary Table S1**, or from previously published RNA-seq libraries listed in **Supplementary Table S4** were mapped to *A. ceylanicum* v2.1 genes by Salmon 1.9.0 (Patro *et al*. 2017).

**Supplementary Table S13.** Statistical analyses of the overlaps between *A. ceylanicum* genes encoding protein motifs (or other traits) and *A. ceylanicum* genes with either intestine-biased or non-intestine-biased gene expression. Statistical significances of overlaps between motif/trait genes and intestine-biased or non-intestine-biased genes were computed and described as in **Supplementary Table S7**. In addition, pairwise overlaps between intestine- or non-intestine-biased gene sets and other gene categories are provided in the ‘Various’ spreadsheet; since these do not involve multiple comparisons of entire motif sets, only p-values rather than q-values are computed for these overlaps. *A. ceylanicum* gene categories in “Various” are: “Intest_haemonchus”, genes with homology to *H. contortus* genes with intestine-biased expression; “Non-intest_haemonchus”, genes with homology to *H. contortus* genes with non-intestine-biased expression; “WashU-intestine-biased”, genes with intestine-biased expression computed from previously published *A. ceylanicum* RNA-seq data; “WashU-non-intestine-biased”, genes with non-intestine-biased expression computed from previously published *A. ceylanicum* RNA-seq data (**Supplementary Table S4**) (Wei *et al*. 2016; Bernot *et al*. 2020).

**Supplementary Table S14**. Statistical analyses of the overlaps between *H. contortus* genes encoding protein motifs (or other traits) and *H. contortus* genes with either intestine-biased or non-intestine-biased gene expression. Statistical significances of overlaps between motif/trait genes and intestine-biased or non-intestine-biased genes were computed and described as in **Supplementary Table S7**.

**Supplementary Table S15.** Statistical analyses of the overlaps between *A. ceylanicum* genes encoding protein motifs (or other traits) and *A. ceylanicum* genes with positively or negatively immunoregulated intestinal or non-intestinal gene expression. Statistical significances of overlaps between motif/trait genes and positively immunoregulated intestinal genes were computed and described as in **Supplementary Table S7**. In addition, pairwise overlaps between immunoregulated gene sets and other categories (such as “Male-biased”) are provided in the “Various” spreadsheet; since these do not involve multiple comparisons of entire motif sets, only p-values rather than q-values are computed for these overlaps.

**Supplementary Table S16.** Statistical analyses of the overlaps between *A. ceylanicum* genes encoding protein motifs (or other traits) and *A. ceylanicum* genes encoding ES proteins that also have positively immunoregulated gene expression in 19-day intestines. Statistical significances of overlaps between motif/trait genes and positively immunoregulated ES genes were computed and described as in **Supplementary Table S7**. In addition, pairwise overlaps between the positively immunoregulated intestinal ES gene set and other categories (such as “Male-biased”) are provided in the “Various” spreadsheet; since these do not involve multiple comparisons of entire motif sets, only p-values rather than q-values are computed for these overlaps.

**Supplementary Figure 1.**
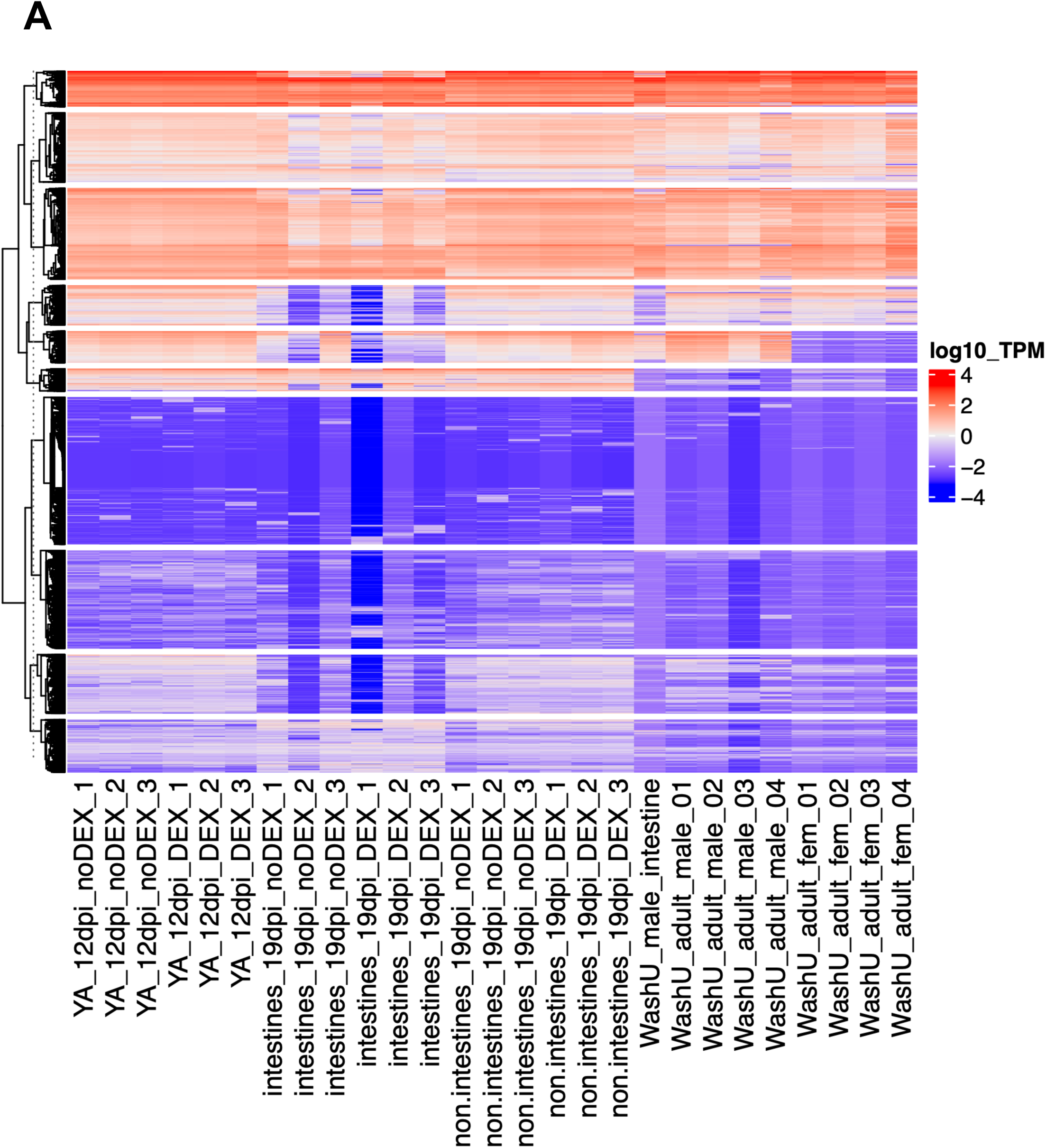
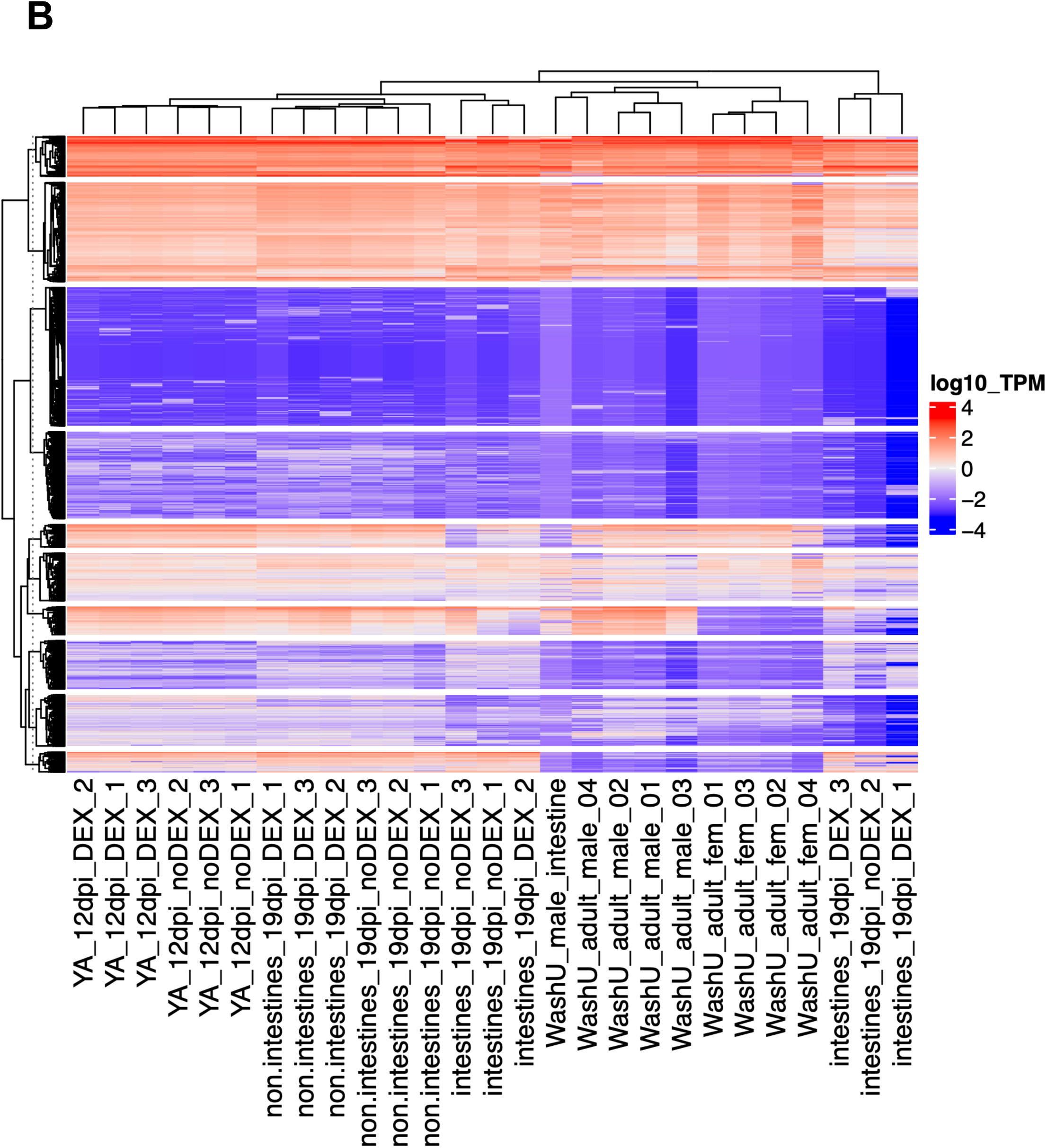
Gene expression in *A. ceylanicum*. Gene activity is shown for all 33,190 *A. ceylanicum* genes, with biological replicates on the x-axis and individual genes on the y-axis. Expression levels are in TPM (log10). In **Supp. Fig. 1A**, replicates are ordered as in Figure 2; in **Supp. Fig. 1B**, replicates are ordered by their similarity of expression (and a dendrogram for their similiaries is shown along the top x-axis). Genes sharing similar patterns of expression are split into 10 clusters. Biological replicates are: young 12-day hookworm adults from normal hosts (YA_12dpi_noDEX); young 12-day adults from immunosuppressed hosts (YA_12dpi_DEX); dissected 19-day intestines from mature hookworm adults in normal hosts (intestines_19dpi_noDEX); dissected 19-day intestines in immunosuppressed hosts (intestines_19dpi_DEX); dissected 19-day intestines in normal hosts (non.intestines_19dpi_noDEX); dissected 19-day non-intestinal tissues in immunosuppressed hosts (non.intestines_19dpi_DEX); previously published adult male intestine (WashU_male_intestine); published adult males (WashU_adult_male); and published adult females (WashU_adult_fem). These heatmaps include two biological replicates (19-day nonDEX intestine replicate 2 and 19-day DEX intestine replicate 2) that clustered anomalously with replicates 1 and 3 of intestinal RNA-seq samples with opposite nonDEX versus DEX status. We interpret this to mean that labeling for the nonDEX/DEX status of these samples was accidentally reversed between RNA harvesting and RNA-seq. We thus omitted these two samples from differential gene expression analysis.

**Supplementary File 1.** An R script used in batch mode for differential gene expression with edgeR 3.36.0 and R 4.1.3.

## Data Availability

*A. ceylanicum* transcriptomic data have been archived in the Sequence Read Archive (Katz *et al*. 2022) with NCBI BioProject accession PRJNA1045065, and with BioSample accessions SAMN38429478, SAMN38440155, SAMN38440168, SAMN38440169, SAMN38440219, and SAMN38440220. Mass spectrometry proteomics data have been deposited to the ProteomeXchange Consortium via the PRIDE partner repository (Perez-Riverol *et al*. 2022) with the dataset identifiers PXD047871 and PXD047879. Protein-coding gene predictions for *A. ceylanicum* have been archived at the Open Science Framework (https://osf.io/dxfsb).

## Acknowledgments

We thank Titus Brown and the Michigan State University High-Performance Computing Center (supported by U.S. Department of Agriculture grant 2010-65205-20361 and NIFA-National Science Foundation (NSF) grant IOS-0923812) for computational support; additional computing was enabled by start-up and research allocations from NSF XSEDE (TG-MCB180039 and TG-MCB190010). We also thank the Millard and Muriel Jacobs Genetics and Genomics Laboratory at the California Institute of Technology for sequencing and computational support.

